# CompleteBin: A transformer-based framework unlocks microbial dark matter through improved short contig binning

**DOI:** 10.1101/2025.04.20.649691

**Authors:** Bohao Zou, Zhenmiao Zhang, Xiaohan Wang, Rong Tao, Nianzhen Gu, Karsten Kristiansen, ShuaiCheng Li, Mo Han, Lu Zhang

## Abstract

Metagenomic binning is crucial for reconstructing microbial genomes from metagenomes. However, existing tools perform poorly in complex communities, where short contigs from low-abundance organisms predominate. This limits the deciphering of community composition and the functions it encodes. Here we present CompleteBin, a Transformer-based framework that bins contigs as short as 850 bp by combining context-based sequence tokenization, taxonomic embeddings from a Transformer pre-trained on reference genomes, and dynamic multi-view contrastive learning.

We benchmarked CompleteBin against five binners using 80 CAMI II and 54 real-world metagenomes, yielding 111 CAMI II evaluation instances through the use of both gold-standard and independently assemblies. AMBER evaluation showed that CompleteBin achieved the highest base-pair-level F1 score in every CAMI II dataset across single-sample, multi-sample and co-assembly modes. CompleteBin also generally recovered more near-complete (NC) and medium-quality-or-higher (MQ+) metagenome-assembled genomes (MAGs) in both CAMI II and the real-world datasets.

In cold-seep sediments, CompleteBin recovered 227 MQ+ strain-level genome bins (StGBs) that no other binner did, spanning 33 phyla. Contigs assigned to MQ+ MAGs only by CompleteBin encoded 473,671 non-redundant genes. Community-scale metabolic modelling further identified 777 StGBs encoding both complete nitrate and sulfate reduction pathways, 154 of which were not fully recovered by other binners, highlighting understudied microbial contributors to local element cycling.

## Introduction

Metagenomic sequencing has transformed our understanding of microbial diversity by enabling culture-independent exploration of microbiomes, revealing vast numbers of previously unknown taxa and their ecological functions [1, 2, 3, 4]. The reconstruction of metagenome-assembled genomes (MAGs) through metagenomic binning, a computational process that clusters assembled contigs derived from the metagenome, represents a cornerstone methodology for linking phylogenetic identity to metabolic potential [5]. The quality of binning results critically influences all downstream analyses, from taxonomic profiling to functional modeling, leading to a demand for robust and efficient binning methods. A remaining challenge in metagenomic binning is the accurate assignment of short contigs (typically *≤*2,500 bp). Recent advances in self-supervised learning have improved metagenomic binning [6, 7], but the persistent difficulty of assigning short contigs continues to limit high-quality MAG reconstruction, particularly in highly diverse communities [8]. The scale of what is lost was recently quantified across 92,187 metagenomes: only about 10% of assembled sequence is captured by medium-quality-or-higher (MQ+, completeness *≥*50%, contamination *≤*10%) MAGs, yet the unbinned remainder harbours the vast majority of the *∼*705,000 bacterial and *∼*27,000 archaeal species-level clades predicted from that dataset [9]. Across diverse metagenomes, we observed that short contigs consistently contributed a growing share of nucleotide bases (**Figure 1a**; R^2^ = 0.67; P = 1.54 *×* 10*^−^*^4^) and complete protein-coding genes (**Figure 1b**; R^2^ = 0.71; P = 3.52 *×* 10*^−^*^4^) as the Nonpareil diversity (Nd) [10] of metagenomes increased. Conversely, the share contributed by long contigs (*>*2,500 bp) showed a negative correlation with Nd of metagenomes (**Figure 1a, b**; **Supplementary Note 1**). Additionally, short contigs were predominantly derived from low-abundance microbial species that face assembly challenges due to insufficient sequencing coverage (one-sided Wilcoxon rank sum test, P *<* 2.2 *×* 10*^−^*^16^ for all datasets, **Figure 1c**; **Supplementary Note 2**), yet these species can play equally important roles in ecosystem functioning [11].

**Figure 1:**
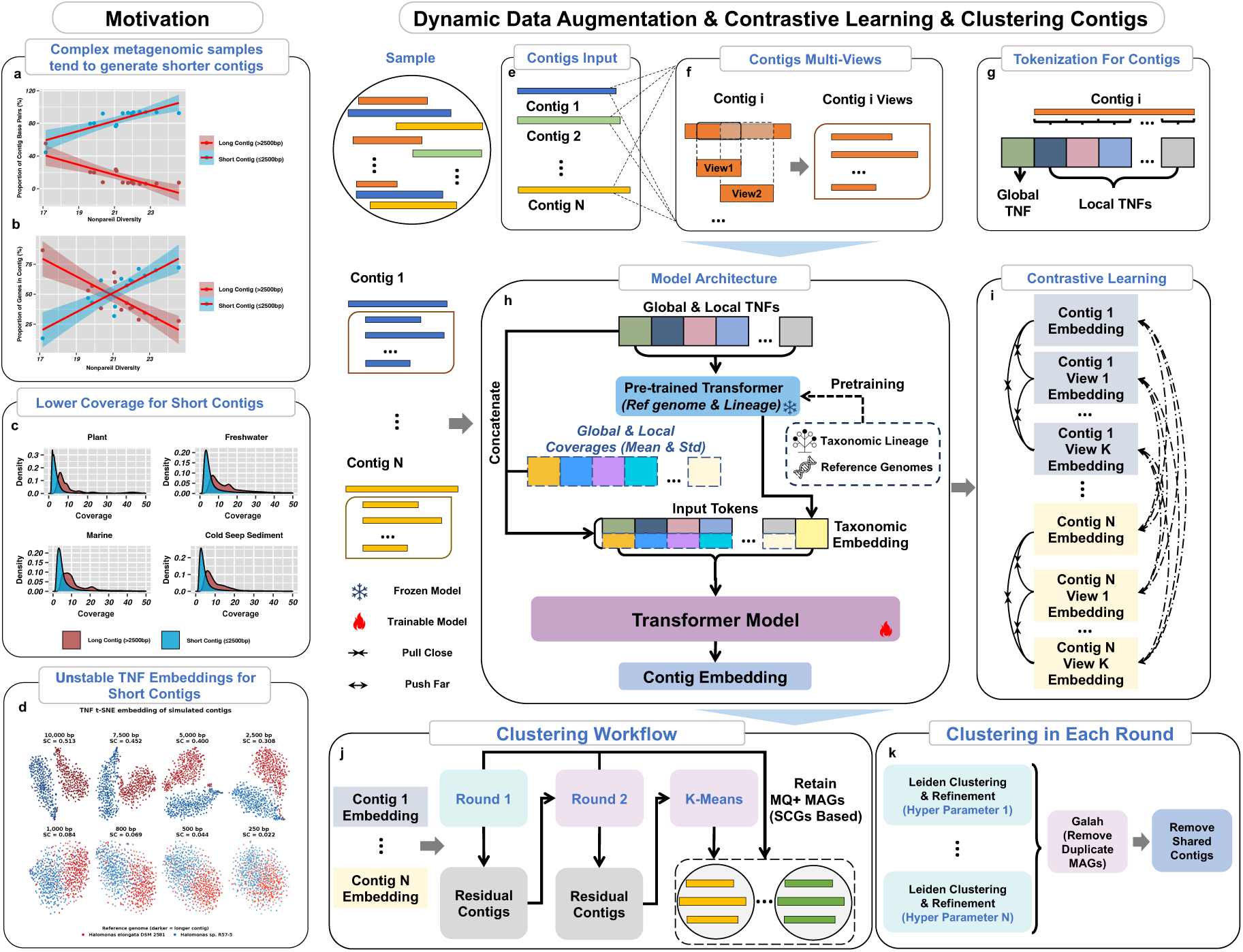
Motivation and framework of CompleteBin. (**a**), The contribution of short contigs (*≤*2,500 bp) to total contig base pairs, plotted against community diversity (Nonpareil diversity index). (**b**), The contribution of short contigs to the number of complete protein-coding genes, plotted against community diversity (Nonpareil diversity index). (**c**), Abundance distributions of short and long contigs in plant, freshwater, marine, and cold seep sediment environments. (**d**), t-SNE embeddings of TNF profiles for simulated contigs of eight lengths (250–10,000 bp) sampled from *Halomonas elongata* DSM 2581 and *Halomonas* sp. R57-5. Colors denote the source genomes, and silhouette coefficients (SC) quantify their separation at each contig length. (**e**), CompleteBin processes a batch with N contigs, and dynamic data augmentation is performed on these contigs at each training step. (**f**), A dynamic data augmentation strategy generates multiple views (subsequences) from each contig within training batches. (**g**), Context-based TNF tokenization is applied for each contig and its augmented views. (**h**), CompleteBin integrates sequence and coverage information via three parallel pathways: (1) a frozen pre-trained Transformer converts context-based TNF tokens into static taxonomic embeddings of contigs; (2) global and local read-depth statistics (mean and standard deviation) are concatenated with the TNF tokens and taxonomic embeddings as input tokens; and (3) a trainable Transformer processes these input tokens to learn dynamic contig representations and generate the final contig embeddings. (**i**), The training objective minimizes distances between a contig and its augmented views (positive pairs) while maximizing distances from other contigs (negative pairs). (**j**), Contigs undergo three rounds of clustering. After each of the first two rounds, only MQ+ MAGs are retained, and contigs not included in the selected MAGs (Residual Contigs) are passed to the next round. The first two rounds use the same clustering method, whereas the final round uses K-means. The MQ+ MAGs retained during the first two rounds and all bins produced in the final round constitute the final output. (**k**), In each of the first two rounds, CompleteBin performs Leiden clustering of the contig embeddings under multiple Leiden-specific hyperparameter settings. It then refines each resulting clustering with single-copy-genes (SCGs). The refined candidate clusterings are subsequently ensembled using Galah. Finally, CompleteBin resolves contigs assigned to multiple MAGs (shared contigs) through an iterative procedure.

Current metagenomic binning tools can be classified into two strategies based on how they process short contigs. The first approach, adopted by MetaBAT2[12], MetaDecoder[13] and TaxVAMB[14], excludes short contigs from MAG reconstruction by default settings. This strategy simplifies the computational problem, but discards all genetic information together with the short contigs that carry it. The second strategy, adopted by MaxBin2[15], CONCOCT[16], SemiBin2[6], and COMEBin[7], attempts to incorporate short contigs (typically 1,000-2,500 bp) into the binning process but relies solely on sequencing coverage and tetranucleotide frequency (TNF), both of which weaken as contigs get shorter. t-SNE embeddings of TNF profiles showed that contigs from two species became difficult to separate as contig length decreased (**Figure 1d**; **Supplementary Note 3**). Additionally, contig sequencing coverage is known to be less effective for single-sample binning [17]. Short contigs are therefore handled poorly by both strategies, which limits genome recovery from complex samples.

To address these challenges, we developed CompleteBin, a deep-learning framework for contig binning that combines three components with an ensemble clustering workflow. 1. Context-Based TNF Tokenization: CompleteBin employs a context-aware tokenization strategy that concatenates global and local TNF vectors into sequential representations. This approach captures both compositional signatures and local sequence context, preserving positional information that becomes particularly important for maintaining specificity in short contigs (**Figure 1g**; **Methods**). 2. Pre-trained Transformer: CompleteBin incorporates external taxonomic knowledge through a Transformer pre-trained on 15,780 publicly available microbial reference genomes and their corresponding species-level taxonomic lineages. The pre-trained model links genomic sequences to taxonomic classifications, effectively compensating for TNF instability in short contigs by leveraging evolutionary relationships learned from comprehensive genomic databases (**Figure 1h**; **Methods**). 3. Dynamic multi-view contrastive learning: to enable high-quality representation learning of contigs with various lengths, CompleteBin uses a contrastive learning framework that dynamically generates multiple subsequence views for each contig during training, and minimizes distances between views derived from the same contig while maximizing distances between views from different contigs (**Figure 1i**; **Methods**). Finally, CompleteBin groups the resulting embeddings with an ensemble clustering workflow that combines Leiden clustering [18] across multiple hyperparameter settings with single-copy-gene-guided (SCG-guided) bin refinement, maximizing genome recovery (**Figure 1j,k; Methods**).

We evaluated CompleteBin by quantifying the contribution of short contigs and benchmarking its performance across diverse metagenomic datasets, including the extremely complex cold-seep sediment communities. CompleteBin in bin-short mode (contigs*≥*850 bp) recovered more MQ+ MAGs for more reference genomes than bin-long (contigs*>*2,500 bp) across all ten CAMI II Marine gold-standard assemblies (**Figure 2a**). Removing short contigs (*≤*2,500 bp) from MAGs generated by the bin-short mode eliminated 71.4-90.6% of additional recoveries across six quality thresholds (completeness *≥* 90%, 70%, 50% and contamination *≤* 5%, 10%), demonstrating their direct contribution (**Figure 2**). We then benchmarked CompleteBin against CONCOCT, MetaBAT2, TaxVAMB, SemiBin2 and COME-Bin using 80 CAMI II [19] and 54 real-world metagenomes. CompleteBin generally recovered more near-complete (NC, completeness *≥*90%, contamination *≤*5%) and MQ+ MAGs across single-sample, multi-sample and co-assembly binning modes and achieved the highest F1 scores in the CAMI II comparisons (**Figures 3–5**, **Supplementary Figures 9 and 11**). In particular, from six cold-seep sediment metagenomes, CompleteBin recovered 227 MQ+ strain-level genome bins (StGBs) spanning 33 phyla that were missed by the other binners and exclusively assigned contigs encoding 473,671 non-redundant genes to MQ+ MAGs, showcasing its strengths in expanding genomic and functional representation of complex microbiomes (**Figure 6**).

**Figure 2:**
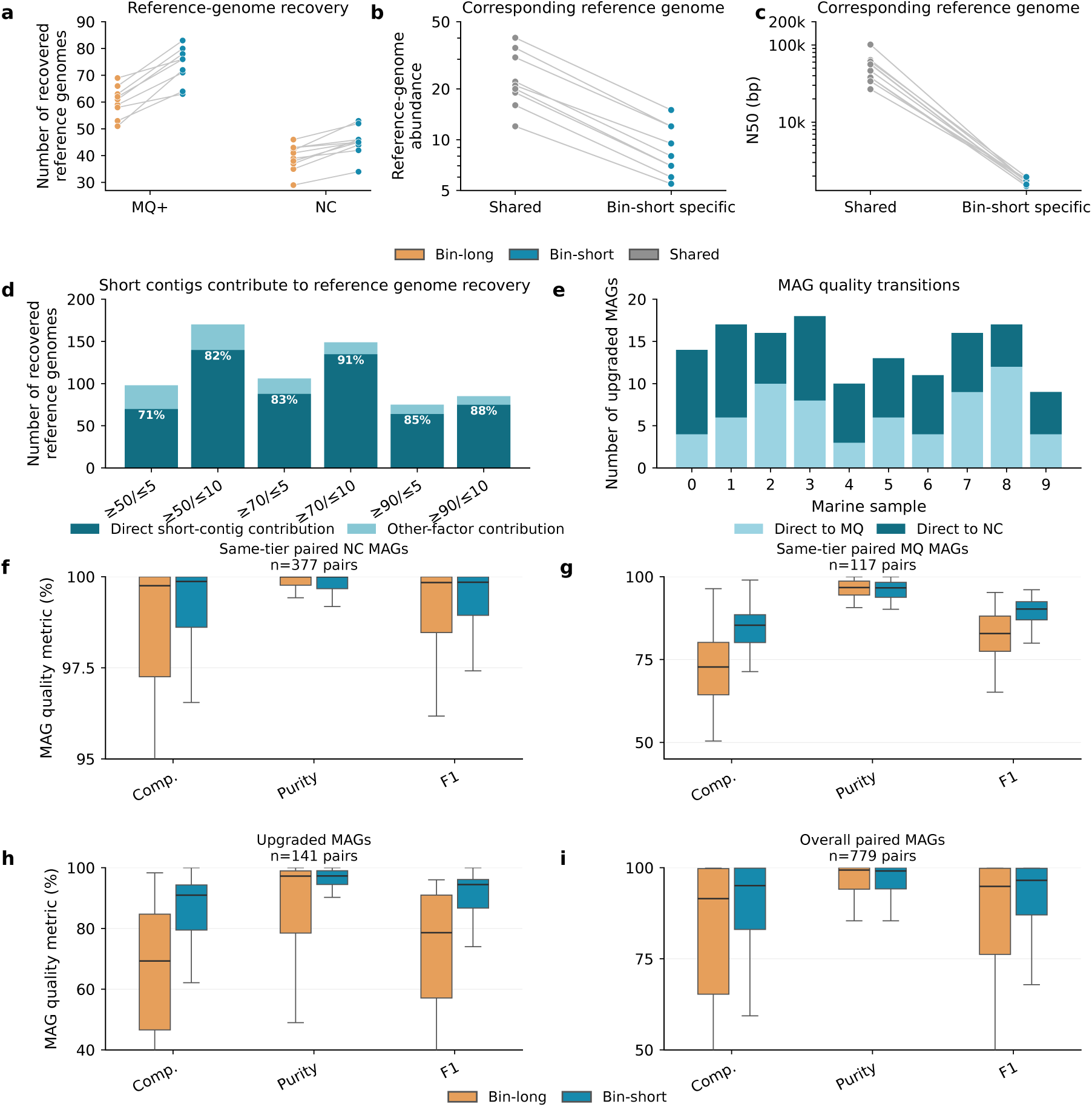
Effects of short contigs on reference-genome recovery and MAG quality across ten CAMI II Marine gold-standard assemblies. (**a**), Paired numbers of reference genomes recovered as medium-quality-or-higher (MQ+) or near-complete (NC) MAGs by bin-long (contigs *>*2,500 bp) and bin-short (contigs *≥* 850 bp); lines connect the same sample (n = 10). (**b**, **c**), Paired sample-level median reference-genome abundance (**b**) and assembly N50 (**c**) for genomes recovered by both configurations (shared) or only by bin-short (bin-short specific). The y axes in b and c are logarithmic, and lines connect groups from the same sample. (**d**), Contributions of short contigs and other workflow factors to reference-genome recovery unique to bin-short under six completeness (*≥*50%, *≥*70% or *≥*90%) and contamination (*≤*5% or *≤*10%) thresholds. Percentages indicate the fraction attributable directly to short contigs. (**e**), Numbers of paired MAGs upgraded to MQ or to NC by bin-short in each sample. (**f** –**i**), Completeness, purity and F1 distributions for same-tier paired NC MAGs (**f**, n = 377 pairs), same-tier paired MQ MAGs (**g**, n = 117 pairs), upgraded MAGs (**h**, n = 141 pairs) and all paired MAGs (**i**, n = 779 pairs).

**Figure 3:**
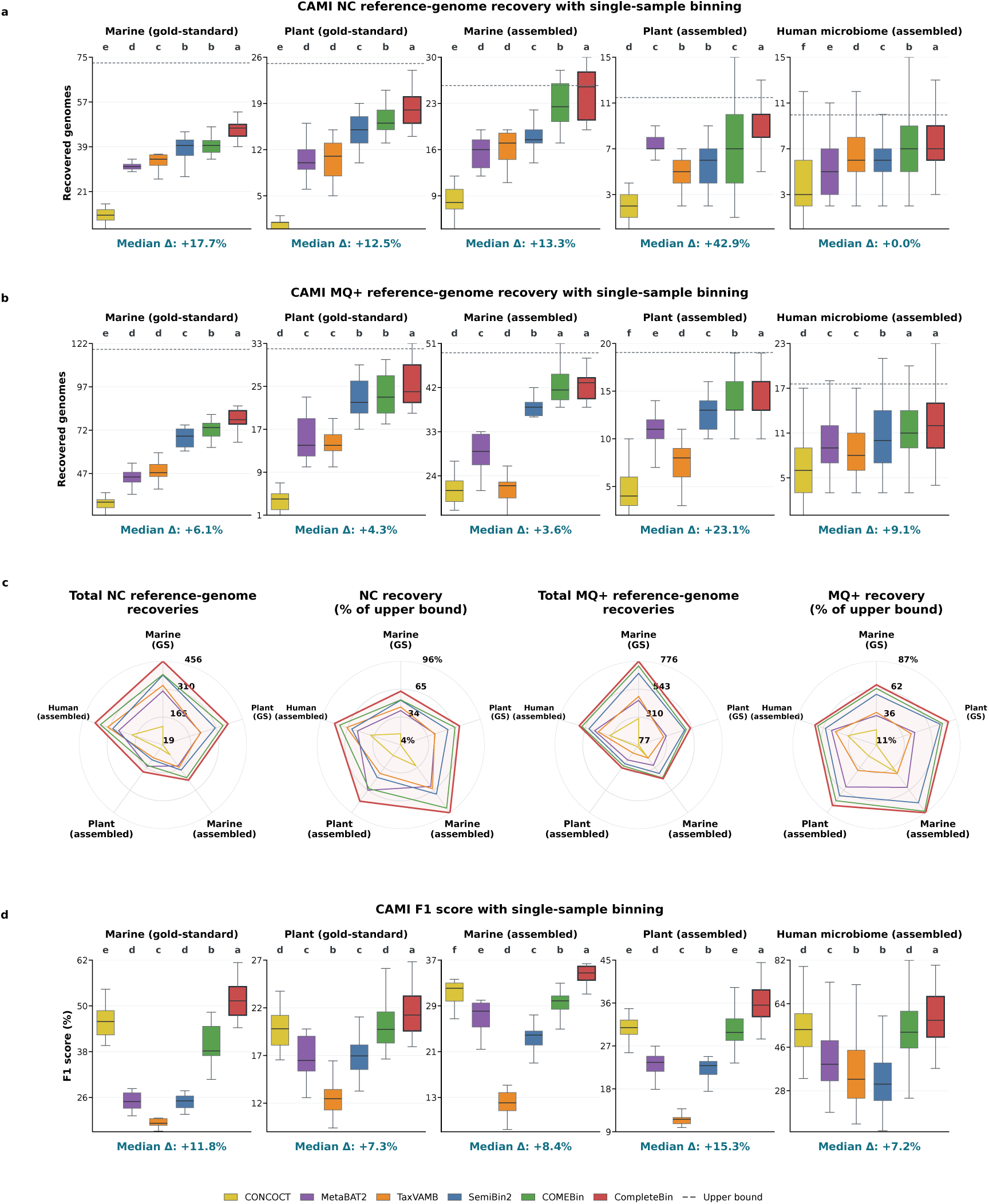
Single-sample binning performance on CAMI II gold-standard and assembled datasets. (**a**, **b**), Numbers of reference genomes recovered as near-complete (**a**; NC) or medium-quality-or-higher (**b**; MQ+) MAGs by six binners across five dataset groups: Marine gold-standard (n = 10), Plant gold-standard (n = 21), Marine assembled (n = 10), Plant assembled (n = 21) and Human microbiome assembled (n = 49). Dashed lines indicate the mean assembly-derived upper bound for each dataset group and quality threshold. (**c**), Radar plots for total NC reference-genome recoveries, NC recovery as a percentage of the corresponding upper bound, total MQ+ reference-genome recoveries and MQ+ recovery as a percentage of the corresponding upper bound across the five dataset groups. (**d**), Base-pair-level F1 scores across the same dataset groups. For CAMI II evaluation, NC recovery requires *≥*90% completeness and *≥*95% purity, whereas MQ+ recovery requires *≥*50% completeness and *≥*90% purity. Compact letter displays group methods that were not significantly different according to unadjusted paired Wilcoxon tests (significance threshold, P *<* 0.05). Median Δ values in (**a, b, d**) report the relative difference between CompleteBin and the second-best binner.

**Figure 4:**
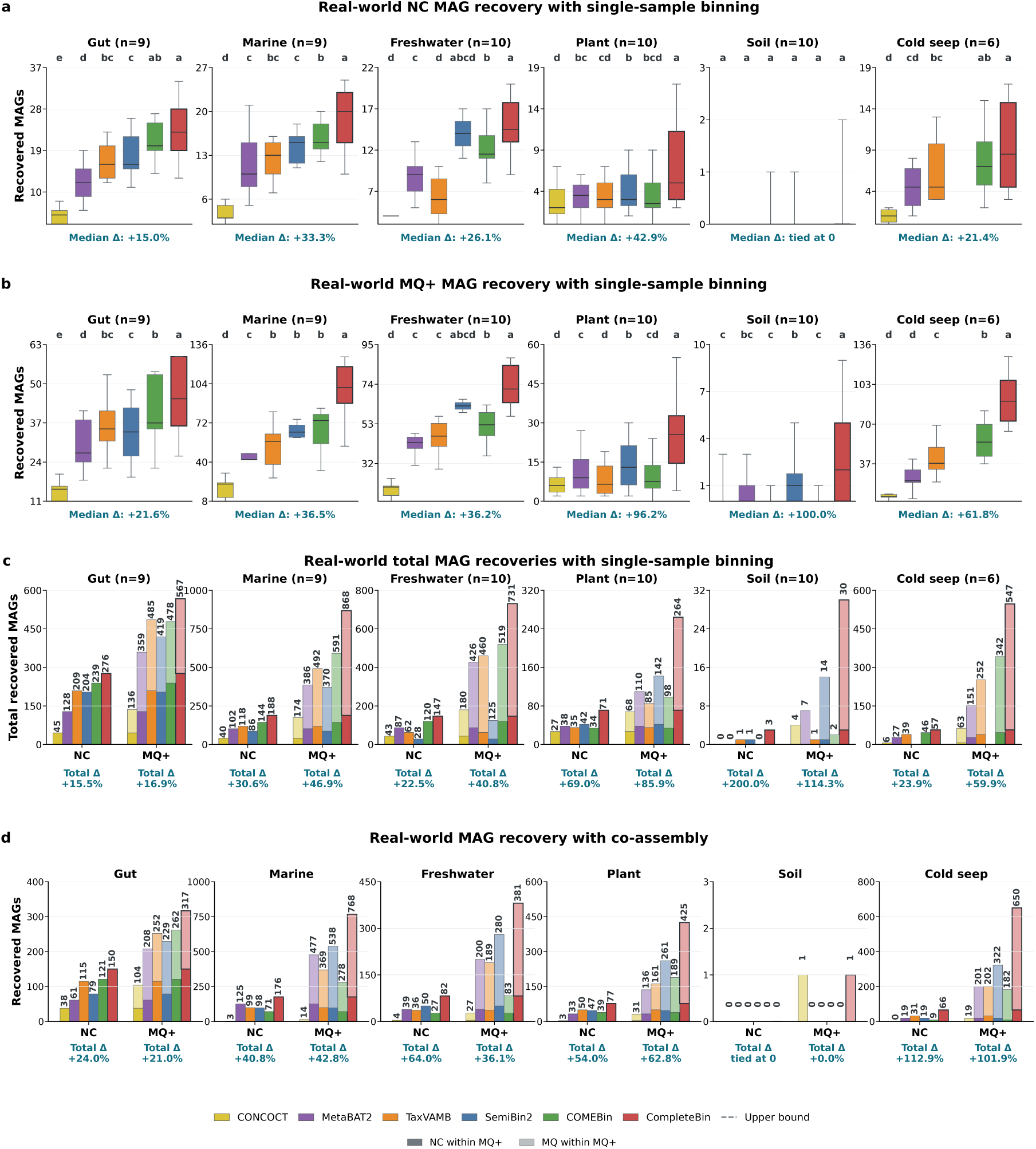
Single-sample and co-assembly binning performance on real-world metagenomes. (**a**, **b**), Numbers of near-complete (**a**; NC) and medium-quality-or-higher (**b**; MQ+) MAGs recovered by six binners in single-sample mode across gut (n = 9), marine (n = 9), freshwater (n = 10), plant (n = 10), soil (n = 10) and cold-seep (n = 6) metagenomes. (**c**), Total NC and MQ+ MAG recoveries obtained by summing the single-sample results within each environment. (**d**), Numbers of NC and MQ+ MAGs recovered from one co-assembly for each environment. NC MAGs have *≥*90% completeness and *≤*5% contamination; MQ+ MAGs have *≥*50% completeness and *≤*10% contamination. Compact letter displays group methods that were not significantly different according to unadjusted paired Wilcoxon tests (significance threshold, P *<* 0.05). Median (**a**, **b**) and total (**c**) Δ values and Δ values for the single co-assembly in (**d**), report the relative difference between CompleteBin and the second-best binner.

**Figure 5:**
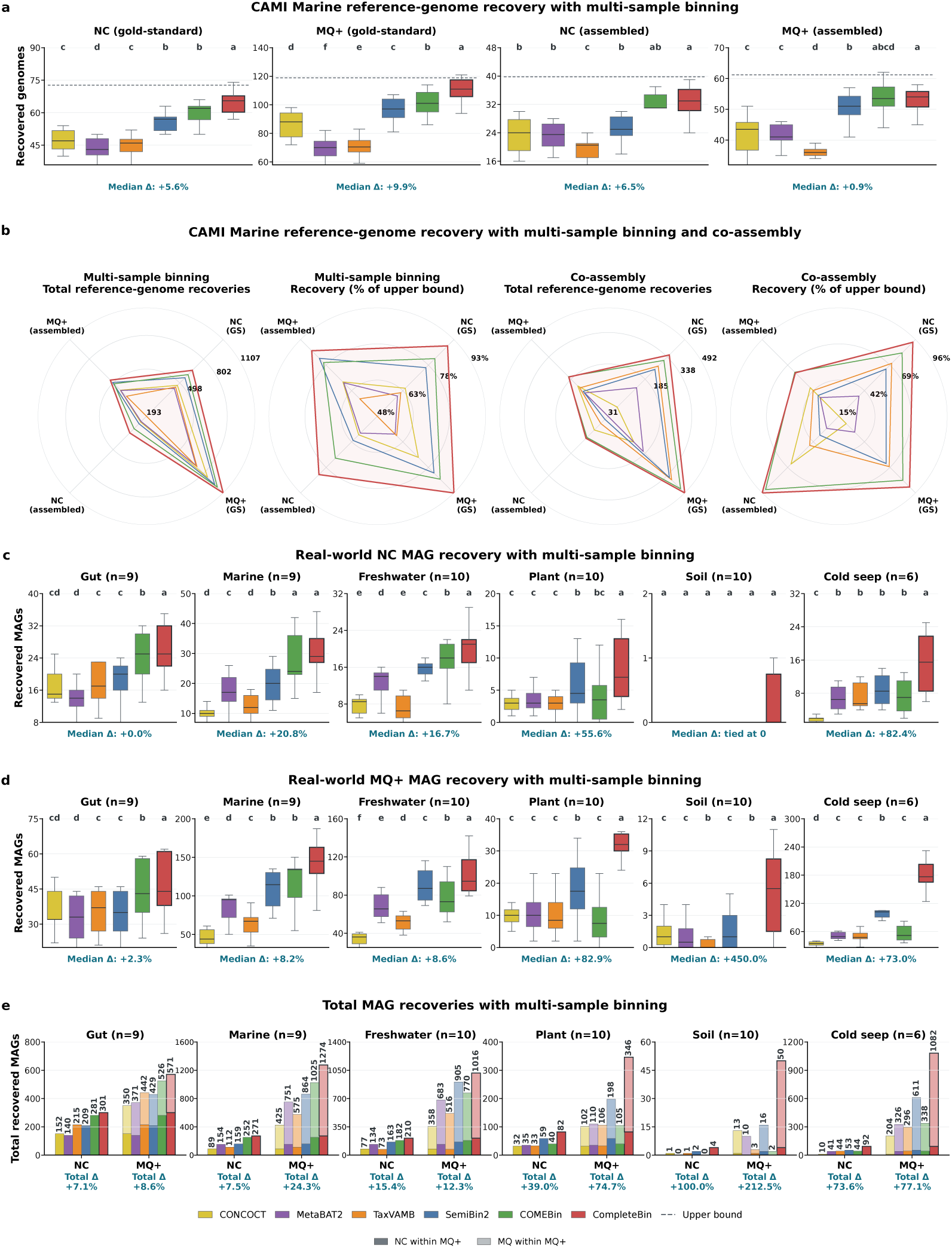
Multi-sample binning performance on CAMI II Marine and real-world metagenomes, and co-assembly binning performance on CAMI II Marine. (**a**), Numbers of CAMI II Marine reference genomes recovered by six binners with multi-sample mode as near-complete (NC) or medium-quality-or-higher (MQ+) MAGs from ten gold-standard and ten metagenome assemblies. Dashed lines indicate the mean assembly-derived upper bounds for the corresponding assembly type and quality threshold. (**b**), Radar plots display total NC and MQ+ reference-genome recoveries from the gold-standard and metagenome assemblies with multi-sample binning. Multi-sample recovery is the percentage of the corresponding upper bound. It also display total NC and MQ+ recoveries from the gold-standard and metagenome co-assemblies, and co-assembly recovery as a percentage of the corresponding upper bound. (**c**, **d**), Numbers of NC (**c**) and MQ+ (**d**) MAGs recovered by multi-sample binning from gut (n = 9), marine (n = 9), freshwater (n = 10), plant (n = 10), soil (n = 10) and cold-seep (n = 6) metagenomes. (**e**), Total NC and MQ+ MAG recoveries obtained by summing the multi-sample results within each real-world environment. Compact letter displays group methods that were not significantly different according to unadjusted paired Wilcoxon tests (significance threshold, P *<* 0.05). Median (**a, c, d**) and total (**e**) Δ values report the relative difference between CompleteBin and the second-best binner.

## Results

### Architecture of CompleteBin

CompleteBin is a Transformer-based framework for binning metagenomic contigs of at least 850 bp. Its representation module comprises context-based TNF tokenization, a pre-trained Transformer, and dynamic multi-view contrastive learning and is followed by an ensemble clustering workflow (**Figure 1e–k**; **Methods**). CompleteBin represents each contig by its sequence composition, coverage profile and a taxonomic embedding learned during pre-training. Context-based TNF tokenization divides a contig into 5 equal-length patches and calculates a local TNF vector for each patch. CompleteBin prepends the global TNF vector of the entire contig to these local TNF vectors, preserving both overall sequence composition and local sequence context (**Figure 1g**). Similarly, CompleteBin calculates coverage features for the entire contig and each contig patch. Each coverage feature comprises the mean and standard deviation of read depth along the nucleotides of the corresponding contig ranges. A Transformer pre-trained on 15,780 representative microbial genomes (**Methods**) and their species-level taxonomic lineages converts the TNF tokens into a taxonomic embedding. CompleteBin concatenates the TNF tokens, coverage features and taxonomic embedding together into a multimodal token sequence, which a trainable Transformer converts into the final contig embedding (**Figure 1h**; **Methods**).

CompleteBin trains the contig embedding with dynamic multi-view contrastive learning. At each training step, CompleteBin generates multiple subsequence views of every contig. The contrastive objective function brings embeddings of the same contig and its views closer while separating them from embeddings of the other contigs in the batch (**Figure 1i**; **Methods**). This training procedure exposes the model to different subsequences throughout training without requiring a fixed set of pregenerated augmentations.

CompleteBin groups the resulting embeddings through three sequential clustering rounds (**Figure 1j**). In the first two rounds, CompleteBin applies Leiden clustering [18] across multiple hyperparameter settings and refines the resulting bins. It then dereplicates its MAGs using Galah [20], and assigns each shared contig to only one MAG using single-copy genes (SCGs; **Figure 1k**; **Methods**). At the end of each of the first two rounds, CompleteBin retains MQ+ MAGs and passes residual contigs to the next round. The third round applies K-means to the remaining contigs, and its bins are combined with the MAGs retained during the first two rounds to generate the final output (**Figure 1j**; **Methods**).

### Short contigs increase genome recovery and MAG quality

Low-abundance microbes suffer from fragmented assemblies due to insufficient sequencing coverage [8], resulting in numerous short contigs (**Supplementary Note 4**). We therefore assessed whether including short contigs improves recovery of low-abundance genomes. CompleteBin was run on contigs of at least 850 bp (bin-short) or longer than 2,500 bp (bin-long) across ten CAMI II Marine gold-standard assembly samples. Reference-genome recovery was evaluated using AMBER at the MQ+ (completeness *≥* 50% and purity *≥* 90%) and NC (completeness *≥* 90% and purity *≥* 95%) thresholds. Because the CAMI II datasets provide gold-standard contig-to-genome assignments, purity is used here in place of the contamination estimates applied to the real-world datasets, and the contamination thresholds are applied as 100%-purity (**Methods**).

Bin-short recovered more reference genomes than bin-long in every sample at both the MQ+ and NC thresholds (two-sided paired Wilcoxon tests, Holm-adjusted P = 0.0039 for both comparisons; **Figure 2a**). We obtained the reference-genome abundances from the official CAMI II abundance profiles. Reference genomes recovered only by bin-short had lower abundance than those recovered by both configurations (one-sided paired Wilcoxon signed-rank test, Holm-adjusted P = 0.0020; **Figure 2b**). They also had lower assembly N50 values, consistent with more fragmented assemblies (one-sided paired Wilcoxon signed-rank test, Holm-adjusted P = 0.0020; **Figure 2c**). These results associated the additional recoveries with low-abundance, fragmented genomes.

We next distinguished the direct contribution of short contigs from the effects of rerunning clustering and post-processing on length-restricted inputs. For each bin-short MAG, we removed contigs of *≤*2,500 bp from its original contig assignments, and recalculated completeness and purity. Of the 170 reference genomes recovered only by bin-short, 140 were no longer recovered because their corresponding MAGs fell below the MQ+ threshold. The remaining 30 MAGs still satisfied the MQ+ threshold after this removal, indicating that their differential recovery was associated with other workflow changes, including training and clustering. Across six combinations of completeness and contamination thresholds, removing short contigs directly accounted for 71.4–90.6% of bin-short-specific recoveries (**Figure 2d**).

We further compared bin-short and bin-long MAGs assigned to the same reference genome. Across the ten samples, 141 bin-short MAGs were assigned to a higher quality tier than their corresponding bin-long MAGs. Of these, 66 moved from low-quality (LQ) to medium-quality (MQ), and 75 moved from LQ or MQ to NC (**Figure 2e**), where MQ denotes MAGs that meet the MQ+ but not the NC threshold (**Methods**). After contigs of *≤*2,500 bp were removed from these 141 bin-short MAGs without rerunning clustering or post-processing, 111 (78.7%) no longer retained their upgraded quality tier, indicating that these quality improvements depended directly on short contigs.

Including more short contigs in a MAG may increase its completeness while reducing purity. To evaluate whether this potential trade-off occurred in CompleteBin, we paired MAGs assigned to the same reference genome across the two configurations, and examined pairs in which both MAGs belonged to the same quality tier (NC, MQ or LQ). The 377 paired NC MAGs had nearly identical completeness, purity and F1 distributions between configurations (**Figure 2f**). Among 117 paired MQ MAGs, bin-short increased mean completeness from 72.1% to 83.3% and mean F1 from 82.0% to 88.9%, while mean purity remained similar (96.5% and 96.0%; **Figure 2g**). Among 106 paired LQ MAGs, bin-short increased mean completeness from 48.7% to 67.2%, mean purity from 61.5% to 67.6%, and mean F1 from 52.0% to 64.5% (**Supplementary Figure 5**). The 141 upgraded MAGs showed a consistent pattern. Relative to their bin-long counterparts, bin-short increased mean completeness from 64.0% to 86.5% and mean F1 from 70.4% to 91.0%. Mean purity also increased from 82.2% to 96.6% (**Figure 2h**). Thus, the completeness gains among paired MAGs were not accompanied by a substantial loss of purity. Across all 779 paired MAGs, which include the 600 same-tier pairs above together with pairs whose two MAGs fell in different tiers, bin-short increased mean completeness from 80.7% to 89.1%, mean purity from 90.7% to 93.5%, and mean F1 from 84.4% to 90.6% (**Figure 2i**). When all MAGs were considered, however, mean purity decreased from 89.0% in bin-long to 78.2% in bin-short. This lower overall mean primarily reflected the larger number of LQ MAGs produced by bin-short (599 versus 229), rather than a loss of purity among paired MAGs (**Supplementary Figure 5**). These results show that short contigs provide the sequence needed to recover low-abundance, fragmented reference genomes and improve the quality of their associated MAGs.

### CompleteBin outperforms state-of-the-art tools in single-sample binning

We benchmarked CompleteBin against five established binners (CONCOCT, MetaBAT2, TaxVAMB, SemiBin2 and COMEBin) using 80 assembled CAMI II metagenomes from the human microbiome (airways (n=10), gastrointestinal (n=10), oral (n=10), skin (n=10) and urogenital (n=9)) and environmental microbiomes (marine (n=10) and plant (n=21)). We also evaluated marine and plant samples using their gold-standard assemblies. We quantified reference-genome recovery by counting the reference genomes represented by at least one recovered MAG meeting the NC or MQ+ threshold (**Methods**). We assessed overall binning quality using base-pair-level completeness, purity and F1 score. For each dataset (assembled and gold-standard assemblies), we estimated the upper bound on the number of reference genomes that could theoretically be recovered as NC and MQ+ MAGs, based on whether enough sequence from each genome was present in the assembly to meet the corresponding completeness threshold (**Methods**). We then divided the observed recovery by this upper bound to quantify the proportion of theoretically recoverable reference genomes that could be generated by each method.

At the sample level, CompleteBin was comparable to or exceeded the second-best method in the median number of reference genomes recovered as NC MAGs across all CAMI II datasets (**Figure 3a**). Relative to the second-best method, the median differences were +17.7% (marine, gold standard), +12.5% (plant, gold standard), +13.3% (marine, assembled) and +42.9% (plant, assembled), with comparable performance in assembled human microbiome samples. At the MQ+ threshold, the corresponding median improvements were +6.1%, +4.3%, +3.6%, +23.1% and +9.1%, respectively (**Figure 3b**). When recoveries were summed across all samples of each dataset, total NC recovery increased by 17.8%, 7.7%, 7.0%, 22.9% and 8.0%, and total MQ+ recovery by 5.3%, 7.5%, 1.9%, 7.2% and 4.0%, respectively (**Figure 3c**, **Supplementary Figure 6**). CompleteBin also achieved the highest F1 score on all datasets, with median improvements of 11.8%, 7.3%, 8.4%, 15.3% and 7.2%, respectively (**Figure 3d**). Across these datasets, CompleteBin ranked first in completeness and placed second to fourth in purity, with TaxVAMB achieving the highest purity on every dataset (**Supplementary Figure 7**). Overall, CompleteBin recovered 63–96% of theoretically recoverable reference genomes at the NC threshold and 65–87% at the MQ+ threshold, consistently matching or exceeding the performance of other binners (**Figure 3c**, **Supplementary Figure 6**). CompleteBin therefore recovered qualifying MAGs for a larger fraction of the reference genomes available in each assembly, while achieving a higher overall binning F1 score.

We next evaluated 54 real-world metagenomes from human gut (n=9), marine (n=9), freshwater (n=10), plant (n=10), soil (n=10) and cold-seep (n=6) communities. Because no gold standard exists for these datasets, NC and MQ+ MAGs were defined using CheckM2 estimates rather than AMBER purity (**Methods**). At the sample level, CompleteBin increased the median number of NC MAGs over the second-best method by 15.0% in gut, 33.3% in marine, 26.1% in freshwater, 42.9% in plant samples and 21.4% in cold-seep samples. All methods recovered a median of zero NC MAGs from soil samples (**Figure 4a**). For MQ+ MAGs, CompleteBin increased the median number by 21.6% in gut, 36.5% in marine, 36.2% in freshwater, 96.2% in plant, 100.0% in soil and 61.8% in cold-seep samples (**Figure 4b**). At the environment level, we summed the numbers of MAGs across all samples within each environment before comparing methods. Relative to the second-best method, CompleteBin increased the total number of NC MAGs by 15.5% in gut, 30.6% in marine, 22.5% in freshwater, 69.0% in plant, 200.0% in soil and 23.9% in cold-seep datasets (**Figure 4c**). The large relative increase in soil is due to the low absolute MAG counts, with three NC MAGs obtained using CompleteBin and one using the second-best method. The total number of MQ+ MAGs increased by 16.9% in gut, 46.9% in marine, 40.8% in freshwater, 85.9% in plant, 114.3% in soil and 59.9% in cold-seep datasets (**Figure 4c**).

### Advantage of CompleteBin persists in multi-sample and co-assembly binning

Multi-sample binning was assessed using ten CAMI II Marine samples and all real-world samples, retaining the independently assembled contigs for each sample while incorporating cross-sample coverage profiles. For co-assembly binning, reads from all samples within each dataset were pooled and assembled jointly, and reads from each sample were then mapped separately to the shared assembly to generate differential coverage profiles (**Methods**).

On the CAMI II Marine dataset in multi-sample binning mode, CompleteBin increased the median reference-genome recovery over the second-best method by 5.6% for NC MAGs and 9.9% for MQ+ MAGs using the gold-standard assemblies. The corresponding improvements using metagenome assemblies were 6.5% and 0.9% (**Figure 5a**). After counts were summed across the ten samples, CompleteBin increased total NC and MQ+ recovery by 9.1% and 9.4%, respectively, for the gold-standard assemblies. Total NC and MQ+ recovery increased by 13.1% and 5.4%, respectively, for the metagenome assemblies. CompleteBin reached 89.4% and 93.1% of the overall NC and MQ+ upper bounds for the gold-standard assemblies (second best: 82.0% and 85.1%). The corresponding values for the metagenome assemblies were 82.4% and 86.6% (second best: 72.9% and 82.2%; **Figure 5b**; **Supplementary Figure 8**). CompleteBin achieved the highest median completeness and F1 score for both assembly types while maintaining median purity above 90% (**Supplementary Figure 9**).

Co-assembly provided a distinct test in which all ten CAMI II Marine samples were assembled jointly. Using the gold-standard co-assembly, CompleteBin represented 355 reference genomes with NC MAGs, 13.8% more than the second-best binner, and 492 with MQ+ MAGs, 8.6% more than the second-best binner. These counts represented 92.7% and 89.3% of the NC and MQ+ upper bounds, respectively (second best: 81.5% and 82.2%; **Figure 5b**; **Supplementary Figure 10**). Using the metagenome co-assembly, CompleteBin represented 107 reference genomes with NC MAGs, 3.9% more than the second-best method, and 198 with MQ+ MAGs, slightly fewer (1.0%) than the best method. These counts represented 95.5% and 60.6% of the corresponding upper bounds (second best: 92.0% and best 61.2%; **Figure 5b**; **Supplementary Figure 10**). CompleteBin achieved the highest F1 score on both co-assemblies (**Supplementary Figure 11**). CompleteBin therefore represented the most reference genomes with NC MAGs and retained the highest F1 score in both co-assembly comparisons.

The advantage extended across the 54 real-world metagenomes in multi-sample mode. At the sample level, CompleteBin tied the second-best median NC MAG count in gut and soil samples, while it increased the median by 20.8% in marine, 16.7% in freshwater, 55.6% in plant, and 82.4% in cold-seep samples (**Figure 5c**). For MQ+ MAGs, the median count increased by 2.3%, 8.2%, 8.6%, 82.9%, 450.0% and 73.0% across gut, marine, freshwater, plant, soil and cold-seep samples, respectively (**Figure 5d**). When MAG counts were summed within each environment, total NC MAG count increased by 7.1%, 7.5%, 15.4%, 39.0%, 100.0% and 73.6%, in the same order. Total MQ+ MAG count increased by 8.6%, 24.3%, 12.3%, 74.7%, 212.5% and 77.1%, respectively (**Figure 5e**). The largest relative gains occurred in plant, soil and cold-seep datasets, particularly at the MQ+ threshold.

We finally compared methods using one real-world co-assembly for each environment. Relative to the second-best method, CompleteBin increased the NC MAG count by 24.0% in gut, 40.8% in marine, 64.0% in freshwater, 54.0% in plant and 112.9% in cold-seep co-assemblies. No method recovered an NC MAG from the soil co-assembly. At the MQ+ threshold, CompleteBin increased MAG counts by 21.0%, 42.8%, 36.1%, 62.8% and 101.9% in gut, marine, freshwater, plant and cold-seep co-assemblies, respectively. In the soil co-assembly, CompleteBin and CONCOCT each recovered one MQ+ MAG and the other methods recovered none (**Figure 4d**). CompleteBin’s advantage therefore persisted when information was shared across samples, through either multi-sample coverage profiles or co-assembly. The largest relative gains were observed in plant and cold-seep co-assemblies.

### Each component of CompleteBin contributes to contig binning performance

To quantify the contribution of each component of CompleteBin, we ablated the three representation-learning components and the ensemble clustering workflow and re-ran binning on three CAMI II Marine gold-standard assemblies (**Supplementary Note 6**). Removing any single representation-learning component reduced the mean F1 score from 0.5137 to between 0.4919 and 0.4929 and lowered the numbers of recovered reference genomes meeting the NC and MQ+ thresholds. Despite producing similar reductions in mean F1 score, the three ablations had distinct effects on completeness, purity and genome recovery. Replacing dynamic subsequence views with fixed views produced the largest reduction in MQ+ genome recovery, recovering 38 fewer MQ+ genomes. Masking the context-dependent tokens produced the lowest mean purity (0.7676). Removing taxonomic pre-training resulted in the loss of 11 NC genomes.

Replacing the ensemble clustering workflow with selection of the best individual Leiden configuration, as in COMEBin, increased mean purity from 0.8033 to 0.8441. However, mean completeness decreased from 0.3777 to 0.3101, and the mean F1 score decreased from 0.5137 to 0.4507, with 17 fewer NC and 40 fewer MQ+ reference genomes recovered. When this ablated configuration was compared with COMEBin, which also selects a single Leiden result, the CompleteBin embeddings yielded higher mean purity (0.8441 versus 0.7647), mean completeness (0.3101 versus 0.2824), and mean F1 score (0.4507 versus 0.4107), and recovered 6 more NC genomes.

### CompleteBin expands the genomic and metabolic representation of cold-seep sediments

CompleteBin recovered 227 MQ+ strain-level genome bins (StGBs) that no other binning tool recovered (**Methods**). Of these, 225 were assigned to 33 phyla, including *Pseudomonadota* (58), *Desulfobacterota* (34) and *Bacteroidota* (17) lineages central to nitrogen and sulfur biogeochemical cycling. Two StGBs could not be classified at the phylum level, which may indicate previously overlooked novel high-level taxonomic groups. Of the 34 *Patescibacteria* StGBs, a candidate phyla radiation group with reduced genomes and metabolic capabilities, eight were recovered only by CompleteBin (**Figure 6a**; **Supplementary Table 1**).

**Figure 6:**
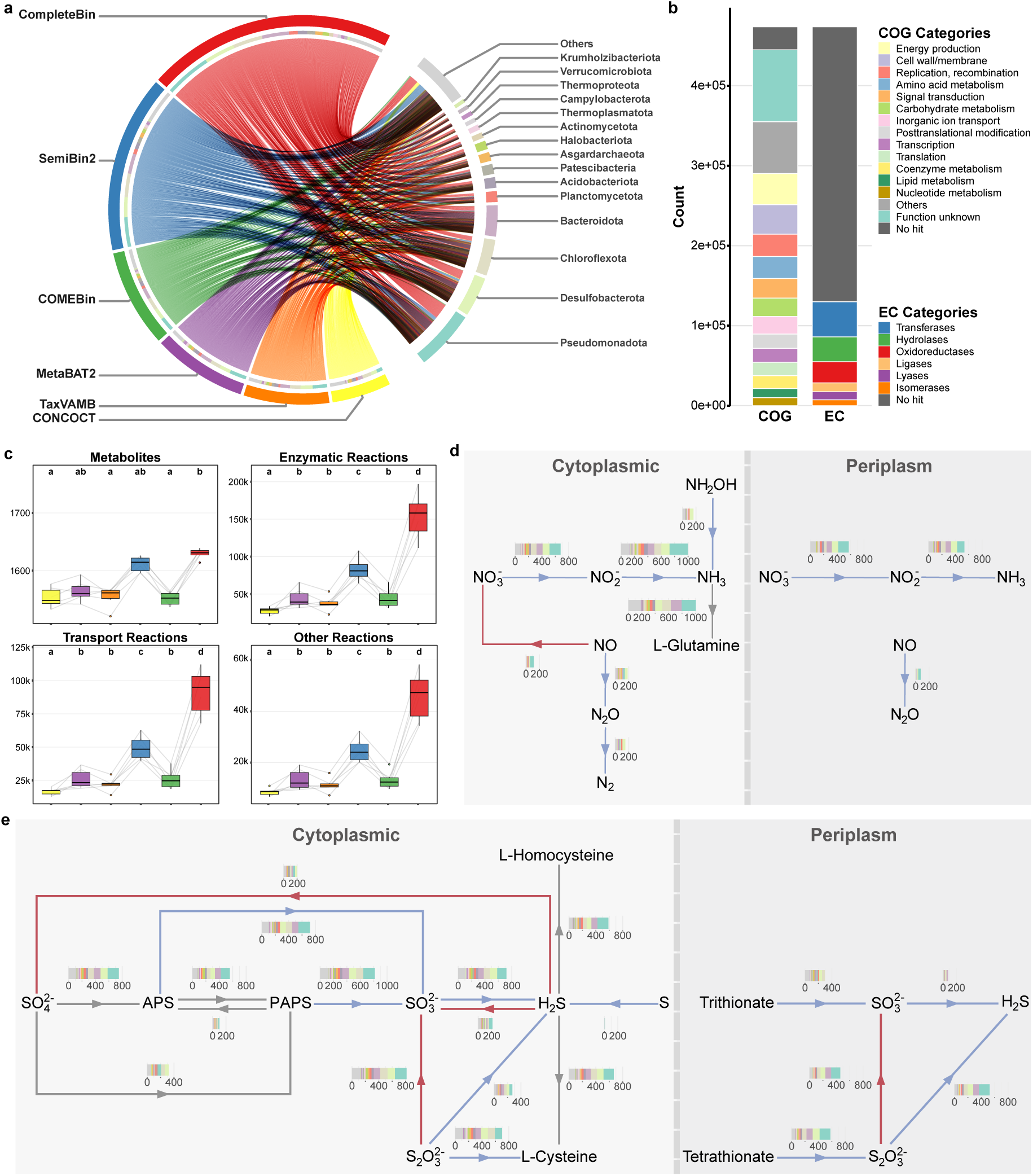
Genomic and metabolic contributions of CompleteBin in cold-seep sediments. (**a**), Medium-quality-or-higher (MQ+) MAGs recovered from cold-seep sediments. The left semicircle shows the six binners, while the right semicircle displays different phyla. Each connecting line between the semicircles indicates that the connected binner on the left recovered a strain-level genome bin (StGB) from the connected phylum on the right. (**b**), Annotation of genes on contigs assigned to MQ+ MAGs by CompleteBin but not by other tools. (**c**), Statistical summary of the community-scale metabolic network. Each sample is considered as an individual community. The four graphs respectively represent the distribution of reactants and products, enzymatic reactions, transport reactions, and other reactions in the network. Box colors denote the six binners as in previous plots. Statistical significance among groups is labeled by compact letter display (Generalized linear mixed model regression followed by Tukey’s honest significant difference post hoc test, *α* = 0.05). (**d**) and (**e**), Metabolic networks of nitrogen (**d**) and sulfur (**e**). Blue lines indicate reduction reactions, and red lines represent oxidation reactions, and grey lines indicate non-redox conversions. Stacked bar charts show the phylum-level distribution of StGBs uniquely identified by CompleteBin as involved in the reactions at corresponding steps, but identified by other tools as not involved.

Comprehensive functional analyses of contigs assigned only to MQ+ MAGs by CompleteBin (**Methods**) recovered 473,671 non-redundant protein encoding genes. Functional categorization identified 172,933 genes related to metabolism, 124,698 involved in cellular processes and signaling, and 69,883 associated with information storage and processing, with 89,927 genes remaining functionally uncharacterized and 28,714 genes lacking COG annotations (note that the functional categories are not mutually exclusive). Only 27.4% of these genes encoded proteins that could be assigned an Enzyme Commission (EC) number, indicating that most fall outside the catalytic functions currently catalogued. Among those with EC assignments, the distribution comprised 44,462 transferases, 26,373 oxidoreductases, 32,970 hydrolases, 11,821 lyases, 8,636 isomerases, and 11,918 ligases (similarly, EC classes are not mutually exclusive; **Figure 6b**), expanding the enzymatic diversity catalogued for this environment.

Community-scale metabolic network modelling assessed the functional significance of CompleteBin’s unique discoveries (**Methods**). While the total number of metabolic substrates changed little, the networks contained more enzymes, transporters and specialized reactions, indicating that the previously unrecovered organisms contribute to metabolism and inter-species interactions in cold-seep sediments (**Figure 6c**). Analysis of nitrogen and sulfur metabolism revealed broad metabolic versatility among the genomes recovered only by CompleteBin. Specifically, the tool identified 236 StGBs with complete nitrate reduction pathways (NO*^−^* to NH_3_, **Figure 6d**) and 227 StGBs capable of complete sulfate reduction (SO^2*−*^ to H_2_S, **Figure 6e**) In contrast, other binners either recovered the corresponding StGBs with the relevant pathways incomplete or did not recover those StGBs at all. Consistent with this pattern, among the 777 StGBs identified by at least one binner as encoding both pathways completely, 154, spanning five phyla (*Desulfobacterota*, *Bacteroidota*, *Pseudomonadota*, *Acidobacteriota* and *Chloroflexota*), were either recovered by binners other than CompleteBin with at least one pathway incomplete, or were not recovered by those binners at all. These 154 dual-pathway organisms suggest previously unrecognized links between nitrogen and sulfur cycling in cold-seep ecosystems, and their potential capacity to use either nitrate or sulfate as an electron acceptor may help explain how sulfate-dependent anaerobic oxidation of methane proceeds under fluctuating geochemical conditions.

## Discussion

Short contigs remain a major obstacle to genome-resolved metagenomics because their composition and coverage signals are sparse, particularly for low-abundance taxa in complex environments. CompleteBin addresses this problem by combining context-based TNF tokenization, dynamic contrastive learning and taxonomic embeddings within a Transformer framework. Unlike approaches based primarily on fixed TNF representations and pre-generated augmentations, this design preserves local and global sequence context across variable contig lengths while exposing the model to diverse sequence views during training. Across CAMI II and real-world datasets, CompleteBin improved MAG completeness and overall binning accuracy, with the largest gains in complex environmental samples, where fragmented assemblies are most prevalent. The compact letter displays in **Figures 3–5** supported this pattern, with CompleteBin in the top statistical group in most comparisons. Better modelling of short contigs is therefore a practical route to recovering genomic information that binning would otherwise discard.

The importance of modelling short contigs becomes particularly evident in complex environmental metagenomes, where assembly fragmentation remains difficult to avoid. A recent analysis [8] showed that soil samples require 0.9-4.6 terabases of clean reads per sample to achieve near-complete metagenomic coverage, over 1,500-fold more than the sequencing depth required for human gut samples. Such depths remain prohibitively expensive with current technologies. Critically, insufficient sequencing coverage directly compromises assembly quality, as evidenced by strong positive correlations between metagenomic depth and both assembly N50 values and read recruitment rates. Even at ultra-deep metagenomic sequencing exceeding 500 Gb, soil metagenome assemblies recruit only 27-57% of reads to contigs longer than 2,000 bp, inevitably yielding substantial proportions of fragmented assemblies dominated by short contigs. For such highly complex environmental microbiomes, assembly fragmentation remains largely unavoidable given practical sequencing and computational constraints. CompleteBin addresses this limitation through context-based TNF tokenization, which preserves compositional information in short sequences, and a pre-trained Transformer whose taxonomic embeddings compensate for the weak compositional signal of short contigs. In cold-seep sediments under multi-sample binning, CompleteBin recovered 73.6% more NC MAGs in total than the second-best method. Improving binning algorithms so that they can exploit short contigs therefore appears more tractable than pursuing the sequencing depth that high-quality *de novo* assembly would require in these environments.

Long-read sequencing presents a promising alternative to short-read technologies for generating high-quality microbial genomes. Several groups now generate long-read soil metagenomes successfully, but the routine application of long-read sequencing to complex environmental samples remains constrained by technical challenges [21]. Key obstacles include the difficulty of obtaining high-quality, high-molecular-weight DNA, the vulnerability of this DNA to damage and contamination [22], and the high cost associated with sequencing low-abundance taxa to adequate coverage in highly diverse communities. Short-read sequencing therefore remains the predominant approach for characterizing complex environmental microbiomes at scale. In our benchmarks, CompleteBin’s advantage was largest in the plant, soil and cold-seep datasets, which are among the most complex and fragmented communities analyzed. This makes it particularly useful for complex environmental samples, for which short-read sequencing remains the most practical option.

CompleteBin requires more computation than COMEBin. Across the seven CAMI II samples examined, the mean wall-clock runtime of CompleteBin with GPU acceleration was 5.0 h, compared with 1.2 h for GPU-accelerated COMEBin (**Supplementary Note 7**). Most of this additional cost arose from the ensemble clustering and post-processing stage, which accounted for 74–81% of the total runtime on the marine samples, whereas model training contributed comparatively less. This overhead reflects a deliberate trade-off: as shown in the ablation analysis, replacing ensemble clustering with a single Leiden configuration reduced completeness and F1 and recovered fewer NC and MQ+ genomes. Peak GPU memory usage did not exceed 11 GB, so training could be performed on a single consumer-grade GPU with sufficient capacity under the tested settings; however, host-memory requirements were substantially higher and increased with input size. Runtime of CompleteBin tended to increase with the Nonpareil diversity of the samples, again driven mainly by clustering and post-processing. Further optimization of the clustering stage, for example through parallelization or by pruning the ensemble of candidate configurations, is therefore expected to yield the largest efficiency gains. This will be particularly important for co-assemblies, which combine contigs from multiple samples into a single, larger binning problem.

A limitation of CompleteBin is its performance on low-complexity datasets with fewer contigs, which provide too few training steps per epoch for stable learning (**Supplementary Note 5**). Such datasets usually arise from high-quality assemblies, from which microbial genomes are comparatively easy to recover.

## Methods

### Overview of CompleteBin

CompleteBin is a Transformer-based deep-learning framework for binning metagenomic contigs. It is designed for contigs of at least 850 bp, with an absolute minimum input length of 768 bp. Its workflow comprises three stages: multimodal contig representation, dynamic contrastive learning and iterative clustering (**Figure 1e–k**). In the first stage, CompleteBin applies context-based tetranucleotide-frequency (TNF) tokenization to each contig and its dynamically generated subsequence views. A frozen, pre-trained Transformer converts the TNF tokens into static taxonomic embeddings. The TNF tokens, taxonomic embeddings and global and local read-depth statistics (means and standard deviations) are concatenated to form a multimodal token sequence. A trainable Transformer processes this sequence to learn dynamic representations and generate the final contig embeddings (**Figure 1f–h**). In the second stage, CompleteBin trains the contig representations using dynamic multi-view contrastive learning. At each training step, the framework generates multiple subsequence views of each contig. The objective brings representations of the same contig and its views closer together while separating them from representations of other contigs in the batch (**Figure 1i**). In the third stage, contigs are processed through three sequential clustering rounds. After each of the first two rounds, MAGs satisfying the MQ+ threshold (which include NC MAGs) are retained. Contigs from the remaining bins proceed to the next round. CompleteBin applies Leiden clustering under multiple hyperparameter settings in these rounds and uses Galah to integrate the resulting bins (**Figure 1k**). Shared contigs across retained bins are then resolved using single-copy gene profiles. The final round applies K-means clustering to the remaining contigs. MAGs retained during the first two rounds and all bins produced in the final round constitute the final output (**Figure 1j**).

### Context-based TNF tokenization and contig coverage features

The TNF vector was generated using a sliding window of size k with a stride of one to traverse the sequence, producing multiple k-mer fragments. The frequency distribution of these k-mer fragments was calculated, resulting in a vector with T dimensions. We set k to 4 and treated reverse complement sequences as equivalent. Consequently, the TNF vector had 136 dimensions (T = 136). The exact formulas used in the calculation are provided below.

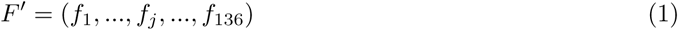

where *f_j_* represents the frequency of the *j*-th k-mer fragment. We further normalized the frequency distribution *F^′^* with

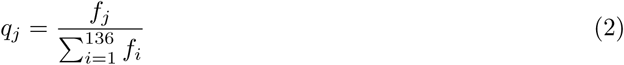

Finally, the TNF vector could be calculated as

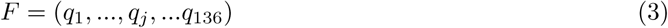

To calculate the local TNF vector, the contig was segmented into a fixed number of patches with equal sequence length. The local TNF vector was calculated separately for each patch to capture local compositional patterns. These local TNF vectors were concatenated in their original sequence order, preserving both local variations and the broader structural context of the sequence. Furthermore, a global TNF vector was added at the beginning of the ordered local TNF vectors. The context-based TNF tokenization for a contig could be calculated as follows:

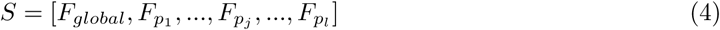

where *F_global_* and *F_p_j__* represent the global TNF vector of the whole contig and the local TNF vector of the *j*-th patch, respectively. *l* is the number of patches. We set *l* to 5 in our study.

We derived global coverage features for each contig following COMEBin. Global features comprised the mean and standard deviation of read depth across all nucleotide positions. Local coverage features for each contig were calculated separately for each individual contig patch using the same statistics as global coverage features. For multi-sample and co-assembly binning, features derived from individual sequencing samples were concatenated into a composite vector for each contig. We then applied the COMEBin normalization procedure to account for differences in sequencing depth and read abundance among samples.

### Architecture of the Transformer in CompleteBin

CompleteBin comprised a frozen, pre-trained Transformer and a trainable Transformer that shared the same backbone architecture. Each model consisted of four Transformer encoder blocks with a hidden dimension of 768. Each block contained a self-attention module followed by a feedforward network [23]. The self-attention module used eight heads, each with a dimension of 96.

The feedforward network had an intermediate dimension of 1,536. It applied a sigmoid linear unit activation [24] to one linear projection. The resulting activations were multiplied element-wise by the output of a second linear projection. A final linear layer projected the gated representation back to 768 dimensions. All linear projections in the self-attention and feedforward modules omitted bias terms.

Root mean square layer normalization [25] was applied before both the self-attention and feedforward sublayers. The query and key representations were also normalized within each attention head before the attention scores were calculated. Learned positional embeddings were added to the token representations to encode token order. During training, an attention dropout probability of 0.15 was applied [26]. Residual connections were retained around both sublayers.

Both Transformers were initialized with the same pre-trained weights. The parameters of the pre-trained Transformer remained fixed, whereas the trainable Transformer was optimized through contrastive learning. The frozen Transformer aggregated its token representations by softmax-weighted pooling to generate a taxonomic embedding. The trainable Transformer used the output representation of the first token as the contig-level representation. A linear projection followed by L_2_ normalization produced the final contig embedding used for clustering.

All model components were implemented in PyTorch v2.8.0, with GPU acceleration provided by CUDA v12.8.

### Pre-training the Transformer model with reference genomes and taxonomic lineages

Reference genomes were obtained from the proGenomes v3.0 database [27], with corresponding complete taxonomic lineages retrieved from the National Center for Biotechnology Information (NCBI). Quality control used a multi-step filtering process. Phyla containing fewer than 10 species were excluded. Redundant entries with identical taxonomy IDs were removed, with preference given to the genome of maximum length. Additionally, we retained only the largest genome per species when multiple strains were present. This filtering procedure yielded a curated dataset of 15,780 representative genomes, each corresponding to a unique microbial species.

To prevent information leakage in the CAMI II benchmarks, CompleteBin used separate pretrained weights for these datasets. We used dRep [28] to compare the 15,780 training genomes against reference genomes from the CAMI II marine, plant-associated and human microbiome datasets. Any training genome sharing at least 95% average nucleotide identity with a CAMI II reference genome was removed. The remaining 15,268 reference genomes were used to train the weights applied in all CAMI II analyses.

A Transformer model with four Transformer encoder layers was employed for pre-training on the curated genomic dataset. The pre-training methodology utilized stochastic sampling of sequences from the reference genomes, with sequence lengths uniformly distributed across a range of 768 to 60,000 base pairs. Input sequences were tokenized using context-based TNF tokenization. The training objective was formulated as a multi-class classification task, wherein each reference genome’s species-level taxonomic annotation constituted a discrete class label. This produced 15,780 classes for the general model and 15,268 classes for the CAMI II-specific model. Model optimization was conducted over 128 training epochs with a batch size of 256. The training procedure employed the AdamW optimizer with an initial learning rate of 1 *×* 10*^−^*^5^, subject to cosine annealing scheduling to facilitate convergence.

### Dynamic contrastive learning framework

#### Dynamic data augmentation

CompleteBin employed the data augmentation strategy during the training process rather than pre-generating augmented data as in SemiBin2 and COMEBin. At each training iteration, a batch of contigs (default batch size: 544) was collected from the dataset, followed by real-time application of data augmentation to generate contrastive pairs. The model was subsequently trained on both the original contigs and their corresponding augmented pairs within the same training step. This procedure was iteratively executed throughout each training epoch. The stochasticity of the data augmentation process would result in identical contigs generating distinct augmented counterparts across different epochs.

#### Dynamic multi-view contrastive learning

CompleteBin dynamically generated multi-views (subsequences) for each contig in a batch during the whole training stage. This optimization goal was designed to pull the representations of a contig and its generated subsequences closer while maintaining the distance between the representations of other contigs and their subsequences within the same batch. The corresponding loss function for two subsequences (*L_ts_*) could be expressed as follows:

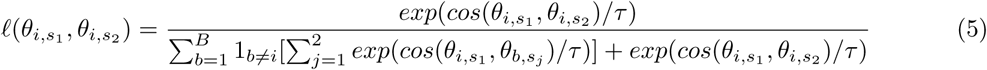

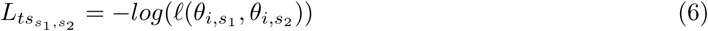

where *θ*_*i,s*_1__ and *θ*_*i,s*_2__ are the representations of the two subsequences of the *i*-th contig. *B* represents the batch size. *cos*(*·, ·*) is the cosine similarity between two vectors, and *τ* is the temperature.

We expanded the formula from two subsequences to multiple subsequences. The final dynamic multi-view contrastive loss (*L*) could be calculated as follows:

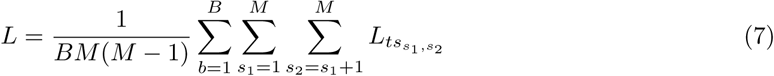

where *M* represents the number of subsequences generated in each training step. During our training process, we set *M* = 6. For each contig, the six contrastive views comprised the original contig and five dynamically sampled contiguous subsequences. New subsequences were generated at every training iteration. The base temperature *τ* was 0.125 for assemblies with N50 *≥* 15,000 bp, 0.145 for those with N50 *≤* 5,000 bp, and 0.135 for intermediate N50 values. The contrastive model was optimized using AdamW with an initial learning rate of 1 *×* 10*^−^*^5^, a weight decay of 1 *×* 10*^−^*^3^, *β*_1_ = 0.9, *β*_2_ = 0.95 and *ɛ* = 1 *×* 10*^−^*^10^. The learning rate was increased tenfold during the first two epochs and subsequently controlled using cosine annealing with warm restarts. The trainable Transformer produced a 768-dimensional representation, which was linearly projected and *L*_2_-normalized to obtain the final 100-dimensional contig embedding used for clustering.

### Clustering

CompleteBin clustered contigs into MAGs through three sequential rounds (**Figure 1j**). The first two rounds combined Leiden clustering [18], single-copy gene (SCG)-based quality assessment and Galah dereplication. After each of the first two rounds, CompleteBin retained MAGs that met the MQ+ threshold based on SCG assessment. Contigs not assigned to retained MAGs were passed to the subsequent round. The third round applied K-means clustering to the remaining contigs. The final output comprised the MQ+ MAGs retained during the first two rounds and all bins generated during the third round.

#### Clustering across multiple graph and Leiden algorithm configurations

In the first round, CompleteBin applied Leiden clustering to normalized contig embeddings. It first constructed k-nearest-neighbour (KNN) graphs following the approach used in COMEBin, with k set to 100. Each graph was sparsified using three edge-retention ratios (the proportions of edges retained for clustering): 50%, 80% and 100%. Following COMEBin, the distance *d_ij_* between connected contigs i and j was converted into an edge weight using the Gaussian-kernel transformation *w_ij_* = exp(*−d_ij_/h*). The bandwidth h controlled how rapidly the edge weight decreased with distance. CompleteBin evaluated bandwidths of 0.05, 0.10, 0.15, 0.20 and 0.25 and Leiden resolution parameters of 1, 3, 5, 8, 10, 15 and 20. Constraint-based clustering assigned contigs carrying the same single-copy gene to different clusters, preserving taxonomic consistency within genome bins. In the second round, CompleteBin applied the same clustering workflow to the normalized embeddings of residual contigs but used a narrower hyperparameter grid. This round evaluated bandwidths of 0.10, 0.15 and 0.20, resolution parameters of 1, 3, 5, 8, 10 and 15, and edge-retention ratios of 50% and 100%.

#### Refinement of highly contaminated Leiden clustering results

Cluster quality was evaluated using the SCG-based approach established in COMEBin. Clusters with contamination above 12% were reclustered using Leiden at resolutions of 1, 3 and 5. The clustering result with the highest score was retained. The score was defined as *score* = ∑ *N_c,t_, c ∈ {*50, 70, 90*}*; *t ∈ {*5, 10*}*, where *N_c,t_* denotes the number of clusters meeting the corresponding completeness (c) and contamination (t) thresholds. Clusters with contamination between 5% and 12% were refined using complementary evidence from SCGs and contig embeddings. Contigs containing SCGs were removed only when their exclusion reduced contamination without causing a loss of completeness greater than 10%. Contigs without SCGs were subsequently evaluated based on their distances from the bin centroid in the embedding space. A contig without SCGs was considered for removal if its centroid distance exceeded three times the mean centroid distance of the other contigs. Clusters with contamination at or below 5% were retained without modification.

#### Ensembling MAGs from Leiden results across hyperparameter settings

CompleteBin generated candidate clustering results by varying the bandwidth for each combination of Leiden resolution and KNN-graph edge-retention ratio. For each parameter combination, CompleteBin selected up to two candidates. The first maximized the number of MAGs with *≥*90% completeness and *≤*5% contamination. The second maximized the number of MAGs with *≥*50% completeness and *≤*10% contamination. After selection across all parameter combinations, CompleteBin pooled the MAGs from every selected candidate and clustered them using Galah [20] at 95% average nucleotide identity. Within each resulting cluster, CompleteBin retained the MAG with the highest SCG-based quality score (*QS* = completeness *−* 4 *×* contamination), yielding a non-redundant set of high-quality MAGs.

#### Removal of shared contigs after Galah processing

Among the MAGs retained after Galah processing, the same contig (a shared contig) could occur in more than one MAG. CompleteBin assigned each shared contig to only one output MAG through an iterative procedure. First, CompleteBin placed the MAGs retained after Galah processing into a candidate MAG set. In the first iteration, SCG completeness and contamination were calculated for each candidate MAG based on its current contig composition. The corresponding SCG-based quality score was then computed. The MAG with the highest quality score was selected and retained all of its contigs. Contigs shared with the selected MAG were removed from all other candidate MAGs, whereas non-shared contigs remained unchanged. The selected MAG was then removed from the candidate set, and any candidate MAG containing no contigs was discarded. In each subsequent iteration, SCG completeness, contamination, and quality scores were recalculated for the remaining candidate MAGs, and the procedure was repeated. The iterative procedure ended when no MAGs remained in the candidate set.

#### K-means clustering of residual contigs

In the third clustering round, CompleteBin applied K-means to contigs that were not assigned to the MQ+ MAGs retained during the first two rounds. The number of clusters, K, was estimated from single-copy gene (SCG) counts among these contigs. For each detected SCG, we counted the number of contigs carrying that gene, and defined K as the rounded 10th percentile of these counts, with a minimum value of three.

### Single-sample, multi-sample, and co-assembly binning

For single-sample binning, metagenomic sequencing reads were subjected to independent assembly to generate sample-specific contigs. Read alignment was performed against the assembled contigs to compute coverage profiles for each contig. The resulting contigs and their corresponding coverage information served as input for the binning procedure.

For multi-sample binning, metagenomic reads from each sample underwent independent assembly to produce sample-specific contigs. Subsequently, reads from all samples were mapped against each sample-specific contig to generate differential coverage profiles across samples. The sample-specific contigs and their cross-sample coverage matrices were then employed as input for multi-sample binning. For co-assembly binning, metagenomic sequencing reads from all samples were jointly assembled to generate a common set of co-assembled contigs. Reads from each sample were then mapped separately to these contigs to calculate differential coverage profiles across samples. The co-assembled contigs and their corresponding cross-sample coverage profiles served as input for co-assembly binning.

### Contribution of short contigs to genome recovery

To quantify how much short contigs contribute to genome recovery, CompleteBin was run twice on each of the ten CAMI II Marine gold-standard assemblies: once on all contigs of at least 850 bp (bin-short) and once on contigs longer than 2,500 bp (bin-long). All other parameters were identical. Both outputs were evaluated with AMBER against the gold-standard contig-to-genome assignments. For each reference genome, we obtained its abundance from the CAMI II abundance profiles and computed the N50 of its gold-standard contigs (assembly N50).

To separate the direct contribution of short contigs from the effect of rerunning training, clustering and post-processing on length-restricted inputs, we took each bin-short MAG, retained its original contig assignments, removed the contigs of *≤*2,500 bp, and recalculated completeness and purity from the remaining contigs without rerunning any part of the workflow. A recovery was attributed directly to short contigs when the corresponding MAG fell below the relevant quality threshold after this removal. The attribution was repeated across six combinations of thresholds, formed by completeness *≥*50%, *≥*70% or *≥*90% crossed with contamination *≤*5% or *≤*10%. MAGs were paired between the two configurations when they were assigned to the same reference genome, and quality tiers (NC, MQ and LQ) were defined as in the Evaluation metrics section, using AMBER completeness and purity.

Statistical comparisons in this analysis used paired Wilcoxon signed-rank tests across the ten samples. Two-sided tests were used for the NC and MQ+ recovery-count comparisons, with Holm correction applied across these two comparisons. Prespecified one-sided tests were used for the directional comparisons of reference-genome abundance and assembly N50, with Holm correction applied across these two comparisons.

### Benchmarking datasets

We evaluated CompleteBin against five established metagenomic binning methods using 80 CAMI II metagenomes and 54 real-world metagenomes. The CAMI II datasets comprised 49 human microbiome samples from five body sites (airways, 10; oral, 10; urogenital, 9; skin, 10; and gastrointestinal tract, 10), 10 marine samples and 21 plant-associated samples. Human microbiome samples were evaluated using metagenome assemblies, whereas the marine and plant samples were evaluated using both metagenome assemblies and CAMI-provided gold-standard assemblies. The CAMI II marine and plant metagenome assemblies were generated with MEGAHIT (v1.2.9) [29]; human microbiome metagenome assemblies were generated with metaSPAdes (v4.2.0) [30]. Reads were aligned to the corresponding assemblies using BWA-MEM (v0.7.17) [31], and the resulting BAM files were sorted using SAMtools (v1.21) [32]. Contig coverage profiles were calculated from the BAM files using PySAM (v0.22.1). Community diversity was quantified for each sample as the Nonpareil diversity index (Nd) [10].

The real-world benchmark comprised 54 metagenomes from six environments: gut (n = 9), marine (n = 9), freshwater (n = 10), plant-associated (n = 10), soil (n = 10) and cold-seep sediment (n = 6). These metagenomes were assembled using MEGAHIT, and reads were processed as described above.

### Evaluation metrics

For CAMI II datasets, binning results were evaluated using AMBER (v2.0.7) [33]. We used AMBER to calculate average base-pair-level purity, completeness, and F1 scores for the complete binning results in the “quality of bins” setting. A reference genome was considered recovered at near-complete quality when at least one assigned MAG had *≥*95% purity and *≥*90% completeness. Recovery at the medium-quality-or-higher threshold required at least one MAG with *≥*90% purity and *≥*50% completeness. For CAMI II, contamination was defined as 100% - purity, so these purity thresholds correspond to the *≤*5% and *≤*10% contamination thresholds applied to the real-world datasets. Assembly-derived upper bounds were calculated separately at the near-complete (NC) and medium-quality-or-higher (MQ+) thresholds. For assemblies with an official gold-standard mapping, contigs were assigned directly to their corresponding reference genomes, only contigs longer than 850 bp were retained for upper-bound estimation. For metagenome assemblies, contigs longer than 850 bp were aligned to the reference genomes using Minimap2 [34], and only alignments with *≥* 95% sequence identity and *≥* 90% query-contig coverage were retained. If a contig had multiple qualifying alignments, it was assigned using the alignment with the highest product of sequence identity and query-contig coverage. For each reference genome, theoretical completeness was calculated as the proportion of its sequence covered by the assigned contigs. Reference genomes with *≤* 100 kbp of assigned contig sequence were excluded because most binning tools do not report bins below this size. Genomes with theoretical completeness of *≥* 90% were counted towards the NC upper bound, whereas those with completeness of *≥* 50% were counted towards the MQ+ upper bound.

For real-world datasets, MAG completeness and contamination were estimated using CheckM2 (v1.0.1) [35]. MAGs with completeness *≥*90% and contamination *≤*5% were classified as near-complete (NC). Medium-quality-or-higher (MQ+) MAGs had completeness *≥*50% and contamination *≤*10%, including all NC MAGs. Performance was summarized using the numbers of NC and MQ+ MAGs recovered from each sample. For analyses requiring mutually exclusive quality tiers, MQ denotes MAGs that meet the MQ+ criteria but not the NC criteria; bins that do not meet the MQ+ criteria were classified as low quality (LQ).

### Experimental configuration for binning algorithms

We compared CompleteBin with CONCOCT, MetaBAT2, TaxVAMB, SemiBin2, and COMEBin. The benchmark environment contained CONCOCT (v1.1.0), MetaBAT2 (v2.18), SemiBin2 (v2.3.0), COMEBin (v1.0.4) and TaxVAMB implemented with VAMB (v5.0.4). The benchmark wrappers used a minimum contig length of 1,000 bp for CONCOCT, SemiBin2, and COMEBin, 1,500 bp for MetaBAT2, and 2,000 bp for TaxVAMB. SemiBin2 was run in its standard configuration with the reclustering step enabled. COMEBin and TaxVAMB were run using their default workflows. Unless explicitly stated, the remaining parameters followed the software defaults.

CompleteBin was implemented in Python and PyTorch. SCGs were identified using Prodigal (v2.6.3) and HMMER (v3.1b2). The default batch size was 544 and the baseline training schedule was 35 epochs. Four epochs were added when the assembly N50 was at least 1,536 bp. For datasets containing fewer than 115,000 training contigs, the number of epochs was increased as the number of contigs decreased, subject to an upper limit of 200 epochs. Neural-network training was performed using one GPU, whereas data processing and clustering used CPU workers.

### Metagenomic analysis of cold-seep sediment samples

Medium-quality-or-higher (MQ+) MAGs obtained from all binning tools were pooled together and then clustered at 99% average nucleotide identity using dRep [28] (v3.5.0) to generate StGBs. Representative MAGs of these StGBs were taxonomically annotated using GTDB-Tk [36] (v2.4.0). Taxonomic annotations of other MAGs within the same clustering group were inherited from their representative MAG.

Gene prediction was conducted on contigs that were exclusively assigned to MQ+ MAGs by CompleteBin using Prodigal [37] (v2.6.3). Predicted genes were clustered using MMseqs2 [38] (v17.b804f) at a 0.9 similarity threshold. Representative genes were annotated using eggNOG-mapper [39] (v2.1.12). Genes assigned to multiple function categories were counted for each of the relevant categories.

For each sample, genome-scale metabolic models were constructed using CarveMe [40] (v1.6.0) for all MQ+ MAGs recovered by each binning tool and were then combined to build a community-level metabolic model for that sample. For each community-level model, a specific substance was counted only once in statistics. Sub-models of nitrogen and sulfur metabolism were respectively extracted from each community-level metabolic network. For each reaction step, all StGBs involved in that step were identified. By integrating all community-level sub-models obtained from all six samples, we identified StGBs capable of performing each step that were detected by at least one binning tool. Among these, StGBs exclusively detected by CompleteBin were subjected to statistical analysis and visualization.

### Statistics

Statistical analyses were performed using R (v4.1.1). Paired comparisons were evaluated using Wilcoxon signed-rank tests and independent group comparisons using Wilcoxon rank-sum tests, both implemented with the wilcox.test function. Where several related comparisons were reported together, P values were adjusted with the Holm method, and figures state whether adjusted or unadjusted values are shown. Compact letter displays in **Figures 3–5** group methods that could not be distinguished by unadjusted paired Wilcoxon tests at P *<* 0.05. Linear regression modelling was conducted using the lm function to assess relationships between continuous variables. To compare metabolic-network properties across binning tools in **Figure 6c**, we fitted generalized linear mixed models (GLMMs) with Binner as a fixed effect and Sample as a random intercept. Post hoc pairwise comparisons were performed using the emmeans package with Tukey’s adjustment (*α* = 0.05).

## Supporting information

The supplementary informations and tables

## Data availability

The CAMI II datasets were obtained from the CAMI II challenge repository (https://cami-challenge.org/datasets/).

The real-world metagenomic sequencing datasets were obtained from public repositories. The freshwater datasets were retrieved from the Genome Sequence Archive at the National Genomics Data Center under run accessions CRR807742, CRR807744, CRR807746, CRR807748, CRR807750, CRR807753, CRR807756, CRR807758, CRR807760 and CRR807762, and were also retrieved from the NCBI Sequence Read Archive (SRA) under accessions SRR26420192, ERR9631077, and ERR4195020. The plant-associated datasets were obtained from the NCBI Sequence Read Archive (SRA) under accessions SRR25582254–SRR25582263 [41] and SRR10968246. The marine datasets were obtained from the SRA under accessions SRR27204692–SRR27204700, and were also acquired from the Joint Genome Institute (JGI) database under project identifiers 1021526, 1125694, and 1102218, as described in [42]. The soil datasets under accessions SRR27942844, SRR27942847, SRR27942848, SRR27942850, SRR27942851, SRR27942852, SRR27942853, SRR27942860, SRR27942866 and SRR27942867.

The gut datasets comprised eight samples from the IBS-D cohort (K0261, K0262, K0265, K0266, K0267, K0268, K0269 and K0271), available from the China National GeneBank (CNGB) under project accession CNP0000334 [43], and one South Korean human-fecal dataset available from the SRA under accession SRR13060973, SRR13060977 [44]. The cold-seep sediment datasets were obtained from CNGB under project accession CNP0005072, with run accessions CNR1069767 (CS01), CNR1069770 (CS03), CNR1069773 (CS05), CNR1069776 (CS08), CNR1069779 (CS10) and CNR1069782 (CS12) [45].

## Code availability

The source code is freely available at https://github.com/ericcombiolab/CompleteBin [46] under an MIT license.

## Acknowledgements

The project is supported by the Young Collaborative Research grant (No. C2004-23Y, No. C2005-24Y), HMRF grant (No. 11221026), and HKBU RCMS (RCMS/24-25/03).

## References

[1] Berg, G. et al. Microbiome definition re-visited: old concepts and new challenges. Microbiome 8, 1–22 (2020).

[2] Bharti, R. & Grimm, D. G. Current challenges and best-practice protocols for microbiome analysis. Briefings in bioinformatics 22, 178–193 (2021).

[3] Quince, C., Walker, A. W., Simpson, J. T., Loman, N. J. & Segata, N. Shotgun metagenomics, from sampling to analysis. Nature biotechnology 35, 833–844 (2017).

[4] Yang, C. et al. A review of computational tools for generating metagenome-assembled genomes from metagenomic sequencing data. Computational and Structural Biotechnology Journal 19, 6301–6314 (2021).

[5] Wang, J. & Jia, H. Metagenome-wide association studies: fine-mining the microbiome. Nature Reviews Microbiology 14, 508–522 (2016).

[6] Pan, S., Zhao, X.-M. & Coelho, L. P. Semibin2: self-supervised contrastive learning leads to better mags for short-and long-read sequencing. Bioinformatics 39, i21–i29 (2023).

[7] Wang, Z. et al. Effective binning of metagenomic contigs using contrastive multi-view representation learning. Nature Communications 15, 585 (2024).

[8] Johansen, P. L., Chatzigiannidou, I., Berzina, L., Kristiansen, K. & Brix, S. Unveiling soil microbial diversity through ultra-deep short-read metagenomic sequencing and co-assembly. Imeta e70075 (2025).

[9] Prasoodanan PK, V. et al. Unbinned contigs expand known diversity in the global microbiome. Nature Microbiology 11, 1437–1449 (2026).

[10] Rodriguez-r, L. M. & Konstantinidis, K. T. Nonpareil: a redundancy-based approach to assess the level of coverage in metagenomic datasets. Bioinformatics 30, 629–635 (2014).

[11] Yorki, S. et al. Comparison of long-and short-read metagenomic assembly for low-abundance species and resistance genes. Briefings in Bioinformatics 24, bbad050 (2023).

[12] Kang, D. D. et al. Metabat 2: an adaptive binning algorithm for robust and efficient genome reconstruction from metagenome assemblies. PeerJ 7, e7359 (2019).

[13] Liu, C.-C. et al. Metadecoder: a novel method for clustering metagenomic contigs. Microbiome 10, 46 (2022).

[14] Kutuzova, S. et al. Improving metagenome binning by integrating intrinsic features and taxonomy. Nature Biotechnology 1–11 (2026).

[15] Wu, Y.-W., Simmons, B. A. & Singer, S. W. Maxbin 2.0: an automated binning algorithm to recover genomes from multiple metagenomic datasets. Bioinformatics 32, 605–607 (2016).

[16] Alneberg, J. et al. Binning metagenomic contigs by coverage and composition. Nature methods 11, 1144–1146 (2014).

[17] Mattock, J. & Watson, M. A comparison of single-coverage and multi-coverage metagenomic binning reveals extensive hidden contamination. Nature Methods 20, 1170–1173 (2023).

[18] Traag, V. A., Waltman, L. & Van Eck, N. J. From louvain to leiden: guaranteeing well-connected communities. Scientific reports 9, 1–12 (2019).

[19] Meyer, F. et al. Critical assessment of metagenome interpretation: the second round of challenges. Nature methods 19, 429–440 (2022).

[20] J., W. B. Galah: More scalable dereplication for metagenome assembled genomes. (Github) (2024).

[21] Burian, J. et al. Bioactive molecules unearthed by terabase-scale long-read sequencing of a soil metagenome. Nature Biotechnology 1–8 (2025).

[22] Kim, C., Pongpanich, M. & Porntaveetus, T. Unraveling metagenomics through long-read sequencing: a comprehensive review. Journal of Translational Medicine 22, 111 (2024).

[23] Vaswani, A. Attention is all you need. Advances in Neural Information Processing Systems (2017).

[24] Elfwing, S., Uchibe, E. & Doya, K. Sigmoid-weighted linear units for neural network function approximation in reinforcement learning. Neural networks 107, 3–11 (2018).

[25] Zhang, B. & Sennrich, R. Root mean square layer normalization. Advances in neural information processing systems 32 (2019).

[26] Srivastava, N., Hinton, G., Krizhevsky, A., Sutskever, I. & Salakhutdinov, R. Dropout: a simple way to prevent neural networks from overfitting. The journal of machine learning research 15, 1929–1958 (2014).

[27] Fullam, A. et al. progenomes3: approaching one million accurately and consistently annotated high-quality prokaryotic genomes. Nucleic acids research 51, D760–D766 (2023).

[28] Olm, M. R., Brown, C. T., Brooks, B. & Banfield, J. F. drep: a tool for fast and accurate genomic comparisons that enables improved genome recovery from metagenomes through de-replication. The ISME journal 11, 2864–2868 (2017).

[29] Li, D., Liu, C.-M., Luo, R., Sadakane, K. & Lam, T.-W. Megahit: an ultra-fast single-node solution for large and complex metagenomics assembly via succinct de bruijn graph. Bioinformatics 31, 1674–1676 (2015).

[30] Nurk, S., Meleshko, D., Korobeynikov, A. & Pevzner, P. A. metaspades: a new versatile metagenomic assembler. Genome research 27, 824–834 (2017).

[31] Li, H. Aligning sequence reads, clone sequences and assembly contigs with bwa-mem. arXiv preprint arXiv:1303.3997 (2013).

[32] Li, H. et al. The sequence alignment/map format and samtools. bioinformatics 25, 2078–2079 (2009).

[33] Meyer, F. et al. AMBER: Assessment of metagenome BinnERs. GigaScience 7, giy069 (2018). URL 10.1093/gigascience/giy069.

[34] Li, H. Minimap2: pairwise alignment for nucleotide sequences. Bioinformatics 34, 3094–3100 (2018).

[35] Chklovski, A., Parks, D. H., Woodcroft, B. J. & Tyson, G. W. Checkm2: a rapid, scalable and accurate tool for assessing microbial genome quality using machine learning. Nature Methods 20, 1203–1212 (2023).

[36] Chaumeil, P.-A., Mussig, A. J., Hugenholtz, P. & Parks, D. H. Gtdb-tk: a toolkit to classify genomes with the genome taxonomy database (2020).

[37] Hyatt, D. et al. Prodigal: prokaryotic gene recognition and translation initiation site identification. BMC bioinformatics 11, 1–11 (2010).

[38] Steinegger, M. & Söding, J. Mmseqs2 enables sensitive protein sequence searching for the analysis of massive data sets. Nature biotechnology 35, 1026–1028 (2017).

[39] Cantalapiedra, C. P., Hernández-Plaza, A., Letunic, I., Bork, P. & Huerta-Cepas, J. eggnogmapper v2: functional annotation, orthology assignments, and domain prediction at the metagenomic scale. Molecular biology and evolution 38, 5825–5829 (2021).

[40] Machado, D., Andrejev, S., Tramontano, M. & Patil, K. R. Fast automated reconstruction of genome-scale metabolic models for microbial species and communities. Nucleic acids research 46, 7542–7553 (2018).

[41] Munõz-Ramírez, Z. Y., et al. Exploring microbial rhizosphere communities in asymptomatic and symptomatic apple trees using amplicon sequencing and shotgun metagenomics. Agronomy 14, 357 (2024).

[42] Duncan, A. et al. Metagenome-assembled genomes of phytoplankton microbiomes from the arctic and atlantic oceans. Microbiome 10, 67 (2022).

[43] Zhao, L. et al. A clostridia-rich microbiota enhances bile acid excretion in diarrhea-predominant irritable bowel syndrome. Journal of Clinical Investigation 130, 438–450 (2020).

[44] Kim, C. Y. et al. Human reference gut microbiome catalog including newly assembled genomes from under-represented asian metagenomes. Genome Medicine 13, 134 (2021).

[45] Wang, X., et al. A scalable practice for deep-sea metagenomic studies (2024). Preprint.

[46] Zou, B. The version 1.0.7 of completebin. Zenodo (2025). URL 10.5281/zenodo.17451857.

