## Supplementary material for "CompleteBin: A transformer-based framework unlocks microbial dark matter through improved short contig binning": The supplementary informations and tables: Supp-Info.pdf

### Supplementary Information

#### 1 Supplementary Note

##### Supplementary Note 1

**Data collection and preprocessing.** Metagenomic datasets were obtained from multiple repositories. Freshwater samples (accession numbers: SRR26420192, ERR9631077, ERR4195020), human fecal samples (SRR13060973, SRR13060977), and one plant sample (SRR10968246) were retrieved from the Sequence Read Archive database. Marine samples (1021526, 1102218, 1125694) were sourced from [1]. An additional human fecal sample (K0268, accession CNP0000334) was obtained from the China National GeneBank (CNGB).

**Assembly and gene prediction.** All metagenomic samples were assembled using metaSPAdes with default parameters. The Nonpareil diversity (Nd) [2] of each sample was calculated with the parameter settings `-T alignment -f fasta`. Complete protein-coding genes within assembled contigs were identified using Prodigal under default configuration.

**Statistical analysis.** Total base pairs and gene counts were calculated separately for short and long contigs. Four linear regression models were constructed using the `lm` function in R to examine relationships between Nd and: (1) summed base pairs in short contigs, (2) summed base pairs in long contigs, (3) number of complete genes in short contigs, and (4) number of complete genes in long contigs.

**Comparative analysis of contig length distributions.** To assess the impact of microbial community complexity on assembly outcomes, we performed a comparative analysis of contig length distributions between metagenomic samples exhibiting distinct levels of microbial diversity. Sample K0268 (Nd = 17.15) was selected as a representative of low-complexity microbial communities, whereas sample ERR9631077 (Nd = 21.07) was designated as a high-complexity representative. The distribution of contig lengths following assembly was visualized for both samples (**Supplementary Figure 1**). To evaluate statistical differences between the contig length distributions of low- and high-complexity samples, we employed the Wilcoxon rank-sum test (one-sided; $P < 2.2e^{-16}$). Our results demonstrated that high-complexity metagenomic samples yielded a greater proportion of short contigs compared to low-complexity samples following assembly, indicating that increased microbial diversity negatively impacts assembly contiguity.

#### Supplementary Note 2

**Sample selection and assembly.** Four representative environmental samples were selected for coverage analysis: one freshwater sample (ERR4195020), one plant sample (SRR10968246), one marine sample (1021526), and one cold-seep sediment sample (CS01, accession CNR1069767). All samples were assembled using metaSPAdes with default parameters.

**Read mapping and coverage calculation.** Reads were aligned to their respective assembled contigs using BWA-MEM. Coverage calculations were performed using PySAM to determine the sequencing depth for each contig. Average coverage values were computed separately for short and long contigs within each dataset.

**Statistical analysis of coverage distributions and results.** Coverage distributions between short and long contigs were compared using the Wilcoxon rank-sum test (one-sided, $\alpha = 0.05$). Statistical analyses were conducted to assess whether significant differences in coverage existed between short and long contigs across all environmental sample types. Analysis revealed that short contigs exhibited significantly lower average coverage compared to long contigs across all tested environmental samples, including plant, freshwater, marine, and cold-seep sediment datasets (Wilcoxon rank-sum test, $P < 2.2 \times 10^{-16}$ for all comparisons; **Figure 1c**).

#### Supplementary Note 3

**Simulation of contigs from reference genomes.** Simulated contigs were generated from two reference genomes: *Halomonas elongata* DSM 2581 and *Halomonas* sp. R57-5, both derived from the CAMI II Marine dataset. Contigs were extracted at eight predefined length thresholds: 10,000 bp, 7,500 bp, 5,000 bp, 2,500 bp, 1,000 bp, 800 bp, 500 bp, and 250 bp. For each length threshold, 1,000 independent simulations were performed to ensure statistical robustness. Contigs were randomly sampled from the reference genomes according to the specified length parameters.

**Dimensionality reduction and clustering analysis.** Tetranucleotide frequency (TNF) features were extracted from all simulated contigs. To visualize the distribution of TNF features across length thresholds, t-distributed stochastic neighbor embedding (t-SNE) [3] was applied to project the high-dimensional TNF vectors into two-dimensional space (**Figure 1d**). Silhouette coefficients were calculated for each length threshold to quantitatively assess the separability of contigs originating from different reference genomes. The relationship between silhouette coefficients and contig length was evaluated using linear regression analysis.

The t-SNE visualization demonstrated that TNF profiles from the two distinct species became progressively less distinguishable as contig length decreased (**Figure 1d**). Quantitative analysis revealed a significant positive correlation between silhouette coefficient and contig length (linear regression, $P = 3.9 \times 10^{-4}$; **Figure 1d**), indicating that TNF-based species discrimination is substantially compromised in shorter contigs.

#### Supplementary Note 4

**Dataset selection and abundance definition.** The CAMI II Marine dataset (10 samples) was used to investigate the relationship between microbial abundance and assembly quality. For each sample, its reference genomes were ranked according to their abundance setting in the simulation procedure in descending order. Reference genomes in the upper 50th percentile were classified as high-abundance genomes, while those in the lower 50th percentile were designated as low-abundance genomes.

**Assembly and contig mapping.** All samples were assembled using MEGAHIT with default parameters. Assembled contigs were aligned to their corresponding reference genomes using Minimap2 to determine their genomic origins. Contigs were subsequently categorized based on whether they originated from high-abundance or low-abundance genomes.

**Statistical analysis of contig length distributions.** Contig length distributions were compared between high-abundance and low-abundance genome categories using the Wilcoxon rank-sum test (one-sided, $\alpha = 0.05$). Contigs originating from high-abundance genomes were significantly longer than those from low-abundance genomes (Wilcoxon rank-sum test, $P < 2.2 \times 10^{-16}$; **Supplementary Figure 2**), confirming that low-abundance microorganisms exhibit more fragmented genome assemblies.

#### Supplementary Note 5

For parts of metagenomes from the CAMI II human microbiome dataset, CompleteBin yielded fewer NC MAGs than COMEBin. To investigate this observation, we compared contig count distributions between samples where CompleteBin outperformed COMEBin and samples where it underperformed, based on the number of NC MAGs recovered. Samples in which CompleteBin produced fewer NC MAGs contained significantly fewer contigs than samples in which CompleteBin produced more NC MAGs (Wilcoxon rank-sum test, two-sided, $P = 3.42 \times 10^{-5}$; **Supplementary Figure 3**). These experiments used the contig-count-dependent epoch adjustment described in Methods. However, this adjustment only partially mitigated the effect of small datasets, and low-contig samples remained challenging for CompleteBin. One possible explanation is that such datasets still provide insufficient training opportunities or limited diversity for effective model optimization. The underperforming group had a median of 8,162 contigs, although this descriptive value should not be interpreted as a definitive performance threshold.

#### Supplementary Note 6

**Ablation experimental design.** We performed single-factor ablation experiments to quantify the contributions of the three representation-learning components of CompleteBin: context-based tetranucleotide-frequency (TNF) tokenization, the taxonomically pre-trained Transformer and dynamic multi-view contrastive learning. The experiments used CAMI II Marine gold-standard assemblies from samples 7–9. The full CompleteBin workflow served as the baseline, and each ablated configuration altered only one component while retaining the remaining training and binning procedures. All configurations used the same default random seed (2048). To prevent information leakage, reference genomes sharing at least 95% average nucleotide identity with any CAMI II reference genome were excluded from the pre-training collection, leaving 15,268 reference genomes for training the CAMI II-specific model.

The context-tokenization ablation masked the context-dependent feature tokens (the five local TNF tokens) while retaining the first global token and the final pre-trained taxonomic token. The pre-training ablation disabled the pre-trained Transformer and used a learned null taxonomic token in place of the pre-trained taxonomic embedding. The dynamic-view ablation generated one set of subsequence views for each training contig and reused these fixed views across epochs instead of regenerating views at every training step. We additionally evaluated the downstream clustering workflow using a selection strategy following COMEBin. This ablation ran Leiden clustering across multiple hyperparameter configurations but retained only the single configuration with the highest aggregate recovery score, rather than ensembling MAGs from multiple clustering results and applying the subsequent CompleteBin clustering workflow. The score was defined as

$$S = N_{50,5} + N_{70,5} + N_{90,5} + N_{50,10} + N_{70,10} + N_{90,10}, \qquad (1)$$

where $N_{c,t}$ denotes the number of MAGs with SCG-estimated completeness $\geq c\%$ and contamination $\leq t\%$.

Genome-binning performance was evaluated with AMBER at the base-pair level. We report the mean purity, completeness and F1 score across the three samples. We also counted recovered reference genomes at six combinations of completeness and contamination (100% − AMBER purity) thresholds: completeness ≥50%, ≥70% or ≥90%, each combined with contamination ≤5% or ≤10%. Recovered-genome counts were summed across the three samples. COMEBin results from the same samples were included as an external comparator for the clustering ablation because both approaches ran Leiden clustering across multiple hyperparameter configurations and selected the best individual result.

**Ablation analysis of the representation-learning and clustering components.** The full CompleteBin configuration achieved a mean purity of 0.8033, mean completeness of 0.3777 and mean F1 score of 0.5137 across the three samples (**Supplementary Note Table 1**). Disabling any one of the three representation-learning components reduced all three mean metrics. Removing taxonomic pre-training reduced mean purity, completeness and F1 score to 0.7726, 0.3610 and 0.4919, respectively. Replacing dynamic subsequence views with fixed views reduced these metrics to 0.7688, 0.3629 and 0.4929, whereas masking the context-dependent tokens reduced them to 0.7676, 0.3628 and 0.4926. Among the three representation-learning ablations, masking the context-dependent tokens produced the lowest mean purity, whereas removing taxonomic pre-training produced the lowest mean F1 score.

Compared with the full CompleteBin workflow, all three representation-learning ablations recovered fewer reference genomes at every quality threshold (**Supplementary Note Table 2**). Removing taxonomic pre-training resulted in 3–28 fewer recovered genomes across the six thresholds. Replacing dynamic views with fixed views resulted in 5–38 fewer genomes, whereas masking the context-dependent tokens resulted in 7–25 fewer genomes. The fixed-view ablation produced the largest losses at the two thresholds based on completeness $\geq$50%: 37 fewer genomes at purity $\geq$95% and 38 fewer genomes at purity $\geq$90%. At completeness $\geq$90%, the reductions were smaller but remained evident. Removing taxonomic pre-training recovered 11 and 3 fewer genomes at purity thresholds of 95% and 90%, respectively. The corresponding losses were 13 and 5 genomes for the fixed-view ablation and 10 and 7 genomes after masking the context-dependent tokens. Thus, each representation-learning component improved genome recovery, although the magnitude of its contribution varied across quality thresholds.

Selecting the best individual Leiden result increased mean purity from 0.8033 to 0.8441 but reduced mean completeness from 0.3777 to 0.3101 and mean F1 score from 0.5137 to 0.4507 (**Supplementary Note Table 1**). It also reduced the recovered-genome count at all six thresholds, with losses ranging from 13 to 40 genomes relative to the full workflow (**Supplementary Note Table 2**). The largest loss occurred at completeness $\geq$50% and purity $\geq$90%, for which 40 fewer genomes were recovered, followed by completeness $\geq$50% and purity $\geq$95%, for which 36 fewer genomes were recovered. Thus, although selecting the best individual Leiden configuration produced purer assignments on average, integrating candidate MAGs across multiple Leiden configurations increased genome recovery and provided a better balance between purity and completeness.

We next compared the clustering ablation with COMEBin to evaluate the contig embeddings under comparable Leiden-based selection strategies. The CompleteBin clustering ablation achieved higher mean purity (0.8441 versus 0.7647), completeness (0.3101 versus 0.2824) and F1 score (0.4507 versus 0.4107) than COMEBin (**Supplementary Note Table 1**). The recovery comparison depended on the quality threshold. At completeness ≥90%, the CompleteBin clustering ablation recovered six more genomes at purity ≥95% (115 versus 109) and nine more genomes at purity ≥90% (122 versus 113). It also recovered one more genome at completeness ≥70% and purity ≥90% (174 versus 173). COMEBin recovered more genomes at the remaining three thresholds: 30 more genomes at completeness ≥50% and purity ≥95% (190 versus 160), 13 more genomes at completeness ≥70% and purity ≥95% (160 versus 147), and 20 more genomes at completeness ≥50% and purity ≥90% (213 versus 193). Under these comparable clustering-selection strategies, the CompleteBin embeddings improved the overall purity–completeness balance and increased the recovery of highly complete (≥90%) genomes, but did not uniformly increase recovery at every quality threshold.

Taken together, these ablations support contributions from context-based TNF tokenization, taxonomic pre-training and dynamic multi-view contrastive learning to CompleteBin's performance on the three evaluated CAMI II Marine samples. The clustering ablation further showed that performance depended on both the learned contig representation and the downstream clustering strategy.

| Configuration | Samples | Mean purity | Mean completeness | Mean F1 |
|---|---|---|---|---|
| COMEBin | 3 | 0.7647 | 0.2824 | 0.4107 |
| CompleteBin (full) | 3 | 0.8033 | 0.3777 | 0.5137 |
| Without taxonomic pre-training | 3 | 0.7726 | 0.3610 | 0.4919 |
| Fixed subsequence views | 3 | 0.7688 | 0.3629 | 0.4929 |
| Masked context-dependent tokens | 3 | 0.7676 | 0.3628 | 0.4926 |
| Select best individual Leiden result | 3 | 0.8441 | 0.3101 | 0.4507 |

**Supplementary Note Table 1: Mean AMBER metrics in the ablation analysis.** Values are arithmetic means of sample-level AMBER results for CAMI II Marine gold-standard samples 7–9.

| Configuration | Comp≥50% Purity≥95% | Comp≥70% Purity≥95% | Comp≥90% Purity≥95% | Comp≥50% Purity≥90% | Comp≥70% Purity≥90% | Comp≥90% Purity≥90% |
|---|---|---|---|---|---|---|
| COMEBin | 190 | 160 | 109 | 213 | 173 | 113 |
| CompleteBin (full) | 196 | 174 | 132 | 233 | 201 | 135 |
| Without taxonomic pre-training | 168 (−28) | 155 (−19) | 121 (−11) | 211 (−22) | 182 (−19) | 132 (−3) |
| Fixed subsequence views | 159 (−37) | 149 (−25) | 119 (−13) | 195 (−38) | 177 (−24) | 130 (−5) |
| Masked context-dependent tokens | 171 (−25) | 157 (−17) | 122 (−10) | 212 (−21) | 181 (−20) | 128 (−7) |
| Select best individual Leiden result | 160 (−36) | 147 (−27) | 115 (−17) | 193 (−40) | 174 (−27) | 122 (−13) |

**Supplementary Note Table 2: Recovered reference genomes in the ablation analysis.** Counts were summed across CAMI II Marine gold-standard samples 7–9. For the four ablated CompleteBin configurations, parenthetical values denote differences from the full CompleteBin configuration. Comp, completeness.

#### Supplementary Note 7

**Experimental design.** We evaluated the computational efficiency of CompleteBin using three CAMI II Marine samples (samples 7–9), two CAMI II gastrointestinal-tract samples (samples 0 and 1) and two CAMI II plant-associated samples (samples 1 and 11). Each method was run with 64 CPU cores and up to 300 GB of host memory. CompleteBin was evaluated in both GPU and CPU training modes; COMEBin was also evaluated in GPU and CPU modes; GPU runs used one NVIDIA H20 GPU. Wall-clock time was divided into data processing, model training, and clustering plus post-processing. For CompleteBin, stage-specific times were obtained from its timing records; for COMEBin, total wall-clock time was measured using GNU `time`, and stage boundaries were recovered from timestamps in the COMEBin log. The GPU and CPU implementations of CompleteBin shared the same data-processing and downstream clustering steps and differed only in the hardware used for model training.

Peak host-memory use was obtained from the `MaxRSS` field in Slurm accounting records for the same seven samples used in the runtime analysis. Peak GPU memory during CompleteBin training was measured using `nvidia-smi`.

**CompleteBin required longer runtime than COMEBin.** GPU acceleration reduced the training time of CompleteBin in all seven samples (**Supplementary Note Table 3**). With GPU acceleration for both methods, CompleteBin required 8.382–10.576 h on the three Marine samples, compared with 1.712–2.175 h for COMEBin, and 1.189–2.434 h on the gastrointestinal-tract and plant-associated samples, compared with 0.341–0.954 h for COMEBin. Across these seven samples, the mean total runtime was 5.011 h for GPU-accelerated CompleteBin and 1.179 h for GPU-accelerated COMEBin. Most of this difference arose from clustering and post-processing, which accounted for 74–81% of the GPU-mode CompleteBin runtime on the Marine samples.

CompleteBin could also be run without a GPU, although CPU-only training increased its total runtime. On the Marine samples, the CompleteBin CPU workflow required 18.512–28.806 h, whereas CPU-based COMEBin required 12.605–16.190 h. On the two plant-associated samples, CPU-only CompleteBin required 7.107–7.132 h, compared with 4.075–4.278 h for COMEBin. We therefore recommend GPU training when suitable hardware is available. CompleteBin used 11 GB of GPU memory in our tests, indicating that a GPU with at least 12 GB of memory is sufficient under the tested settings. Training can also be performed on an 11-GB consumer GPU such as the NVIDIA GeForce RTX 2080 Ti, although we recommend at least 12 GB to provide additional memory headroom.

**Runtime increased with microbial diversity.** We next examined the relationship between runtime and microbial diversity across the seven benchmark samples, using Nonpareil diversity (Nd) [2] as the diversity measure. For GPU-accelerated CompleteBin, CPU-only CompleteBin and CPU-based COMEBin, higher-diversity samples required longer runtimes for data processing, training, clustering plus post-processing and the complete workflow (**Supplementary Figure 4**; linear regression, all $P < 0.05$). Total runtime showed a strong linear relationship with diversity for GPU-accelerated

CompleteBin, CPU-only CompleteBin and CPU-based COMEBin ($R^2$ = 0.888, 0.898 and 0.913, respectively). GPU-accelerated CompleteBin also remained faster than the two CPU-based workflows across samples with different diversity levels.

When the individual workflow stages were examined, GPU acceleration substantially reduced CompleteBin training time, particularly for the higher-diversity samples. For GPU-accelerated CompleteBin, clustering accounted for most of the runtime increase associated with diversity. Data processing and model training increased more slowly. These results indicate that GPU acceleration limited the effect of sample diversity on CompleteBin training time, while the downstream clustering workflow became the main runtime cost.

**Peak memory use across the seven runtime-benchmark samples.** Across the same seven samples used for the runtime analysis, CompleteBin's peak host-memory use ranged from 40.9 to 121.6 GiB, whereas that of COMEBin ranged from 24.5 to 65.6 GiB (**Supplementary Note Table 4**). CompleteBin reached its highest values on the three Marine samples (115.3–121.6 GiB). The two gastrointestinal-tract samples peaked at 40.9 and 53.7 GiB. For the two plant-associated samples, CompleteBin's peak host-memory use was 88.7 GiB. The COMEBin peaks were 24.5–41.0 GiB for the gastrointestinal-tract samples and 49.9–52.9 GiB for the plant-associated samples. The corresponding COMEBin peaks for the three Marine samples were 57.9–65.6 GiB.

CompleteBin therefore used more host memory than COMEBin in this analysis. However, its GPU-memory requirement remained low, with a maximum observed use of 11 GB. We recommend GPU training when suitable hardware is available because it substantially reduced runtime without requiring a data-centre-class GPU. A GPU with at least 12 GB of memory provides sufficient capacity and additional memory headroom under the tested settings. Training can also be performed on an 11-GB consumer GPU such as the NVIDIA GeForce RTX 2080 Ti, although the available margin is smaller.

| Dataset | Sample | Method | Data processing (h) | Training (h) | Clustering + post-processing (h) | Total (h) |
|---|---|---|---|---|---|---|
| Marine | marine-7 | CompleteBin (GPU) | 0.136 | 2.210 | 6.629 | 8.975 |
| Marine | marine-7 | CompleteBin (CPU) | 0.136 | 18.870 | 6.629 | 25.635 |
| Marine | marine-7 | COMEBin (GPU) | 0.375 | 1.075 | 0.389 | 1.839 |
| Marine | marine-7 | COMEBin (CPU) | 0.398 | 11.803 | 0.404 | 12.605 |
| Marine | marine-8 | CompleteBin (GPU) | 0.182 | 2.420 | 7.974 | 10.576 |
| Marine | marine-8 | CompleteBin (CPU) | 0.182 | 20.650 | 7.974 | 28.806 |
| Marine | marine-8 | COMEBin (GPU) | 0.412 | 1.413 | 0.349 | 2.175 |
| Marine | marine-8 | COMEBin (CPU) | 0.434 | 15.301 | 0.455 | 16.190 |
| Marine | marine-9 | CompleteBin (GPU) | 0.221 | 1.340 | 6.821 | 8.382 |
| Marine | marine-9 | CompleteBin (CPU) | 0.221 | 11.470 | 6.821 | 18.512 |
| Marine | marine-9 | COMEBin (GPU) | 0.394 | 0.952 | 0.366 | 1.712 |
| Marine | marine-9 | COMEBin (CPU) | 0.401 | 12.147 | 0.413 | 12.961 |
| Gut | sample-0 | CompleteBin (GPU) | 0.022 | 0.969 | 0.198 | 1.189 |
| Gut | sample-0 | CompleteBin (CPU) | 0.022 | 1.456 | 0.198 | 1.676 |
| Gut | sample-0 | COMEBin (GPU) | 0.135 | 0.146 | 0.060 | 0.341 |
| Gut | sample-0 | COMEBin (CPU) | 0.130 | 1.317 | 0.048 | 1.496 |
| Gut | sample-1 | CompleteBin (GPU) | 0.024 | 1.073 | 0.340 | 1.437 |
| Gut | sample-1 | CompleteBin (CPU) | 0.024 | 1.773 | 0.340 | 2.137 |
| Gut | sample-1 | COMEBin (GPU) | 0.182 | 0.242 | 0.093 | 0.517 |
| Gut | sample-1 | COMEBin (CPU) | 0.158 | 2.416 | 0.076 | 2.650 |
| Plant | plant-GS-1 | CompleteBin (GPU) | 0.146 | 0.926 | 1.010 | 2.082 |
| Plant | plant-GS-1 | CompleteBin (CPU) | 0.146 | 5.976 | 1.010 | 7.132 |
| Plant | plant-GS-1 | COMEBin (GPU) | 0.282 | 0.370 | 0.302 | 0.954 |
| Plant | plant-GS-1 | COMEBin (CPU) | 0.262 | 3.824 | 0.193 | 4.278 |
| Plant | plant-GS-11 | CompleteBin (GPU) | 0.157 | 1.048 | 1.228 | 2.434 |
| Plant | plant-GS-11 | CompleteBin (CPU) | 0.157 | 5.721 | 1.228 | 7.107 |
| Plant | plant-GS-11 | COMEBin (GPU) | 0.243 | 0.325 | 0.147 | 0.714 |
| Plant | plant-GS-11 | COMEBin (CPU) | 0.241 | 3.697 | 0.137 | 4.075 |

**Supplementary Note Table 3: Stage-specific wall-clock runtime of CompleteBin and COMEBin.** All values are in hours.

| Dataset | Sample | CompleteBin peak host memory (GiB) | COMEBin peak host memory (GiB) |
|---|---|---|---|
| Marine | marine-7 | 121.6 | 59.7 |
| Marine | marine-8 | 115.3 | 65.6 |
| Marine | marine-9 | 121.1 | 57.9 |
| Gut | sample-0 | 40.9 | 24.5 |
| Gut | sample-1 | 53.7 | 41.0 |
| Plant | plant-GS-1 | 88.7 | 52.9 |
| Plant | plant-GS-11 | 88.7 | 49.9 |

**Supplementary Note Table 4: Peak host-memory use for the seven samples included in the runtime benchmark.**

#### 2 Supplementary Figures

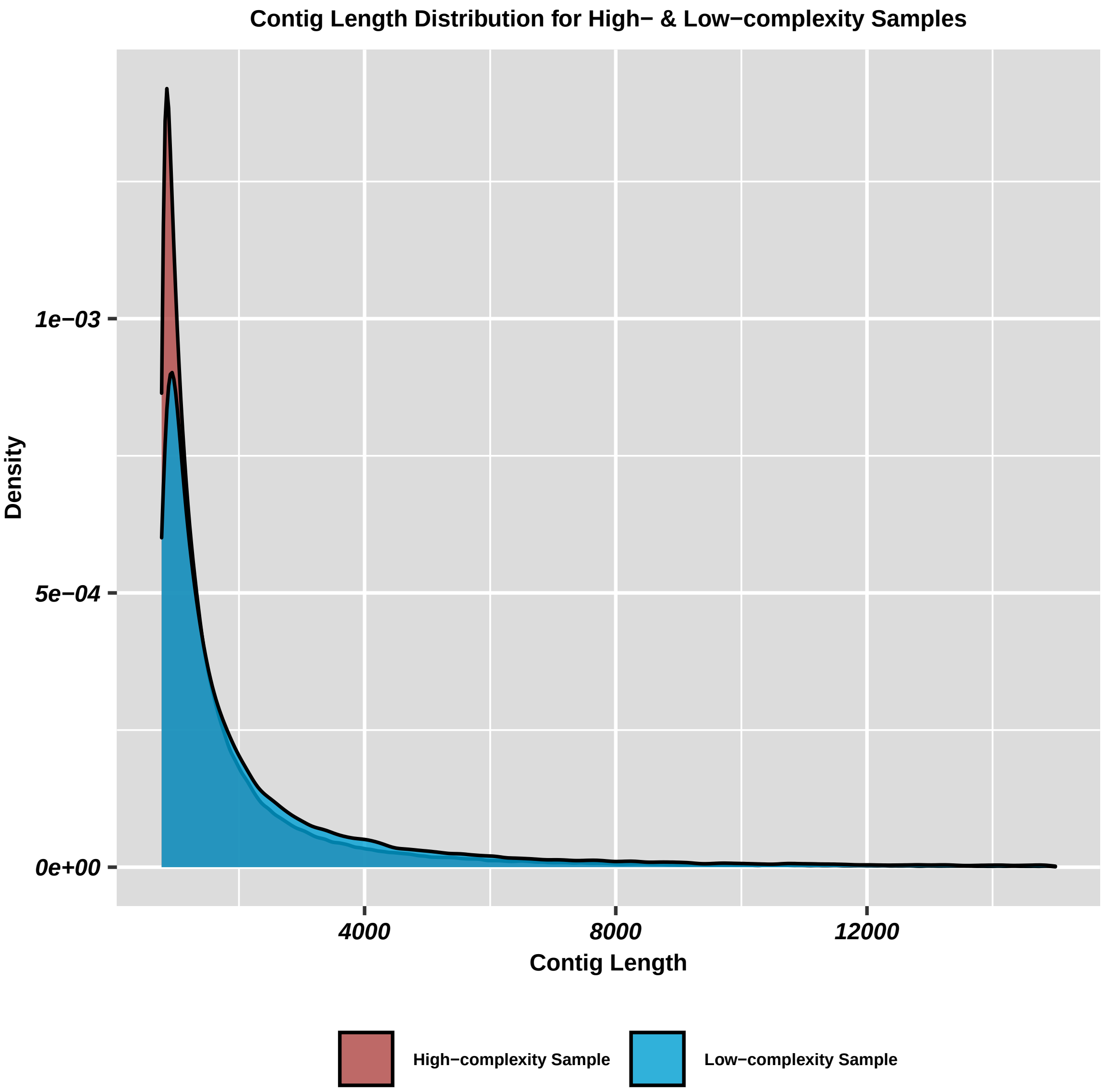


**Supplementary Figure 1: The distribution of contig length after assembly for a high-complexity sample (ERR9631077; Nd = 21.07) and a low-complexity sample (K0268; Nd = 17.15).**

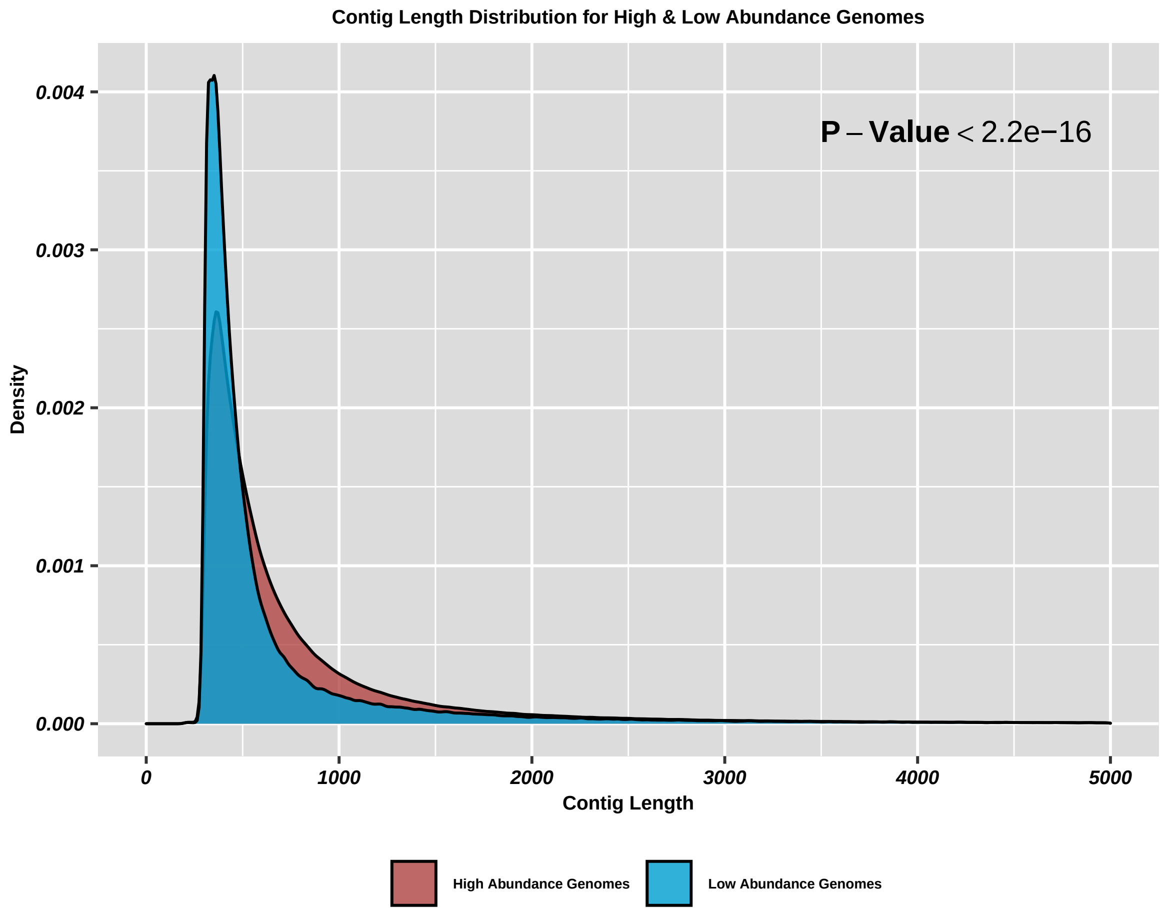


**Supplementary Figure 2: The contig length distribution for the high- and low-abundance reference genomes after assembly.**

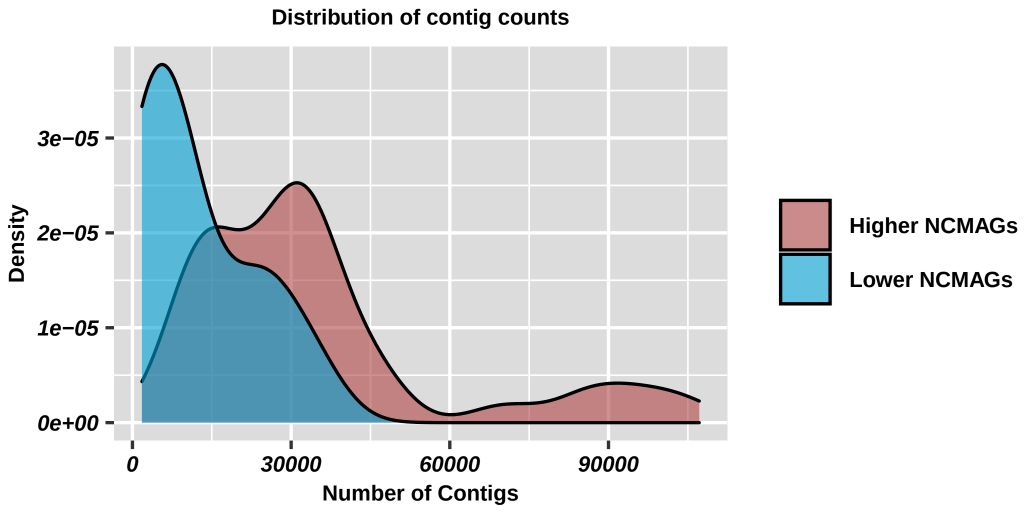


**Supplementary Figure 3: Contig count distribution for metagenomic samples in the CAMI II human microbiome dataset after assembly.** Red indicates samples where CompleteBin yielded more near-complete (NC) MAGs than COMEBin; blue indicates samples where CompleteBin yielded fewer NC MAGs than COMEBin.

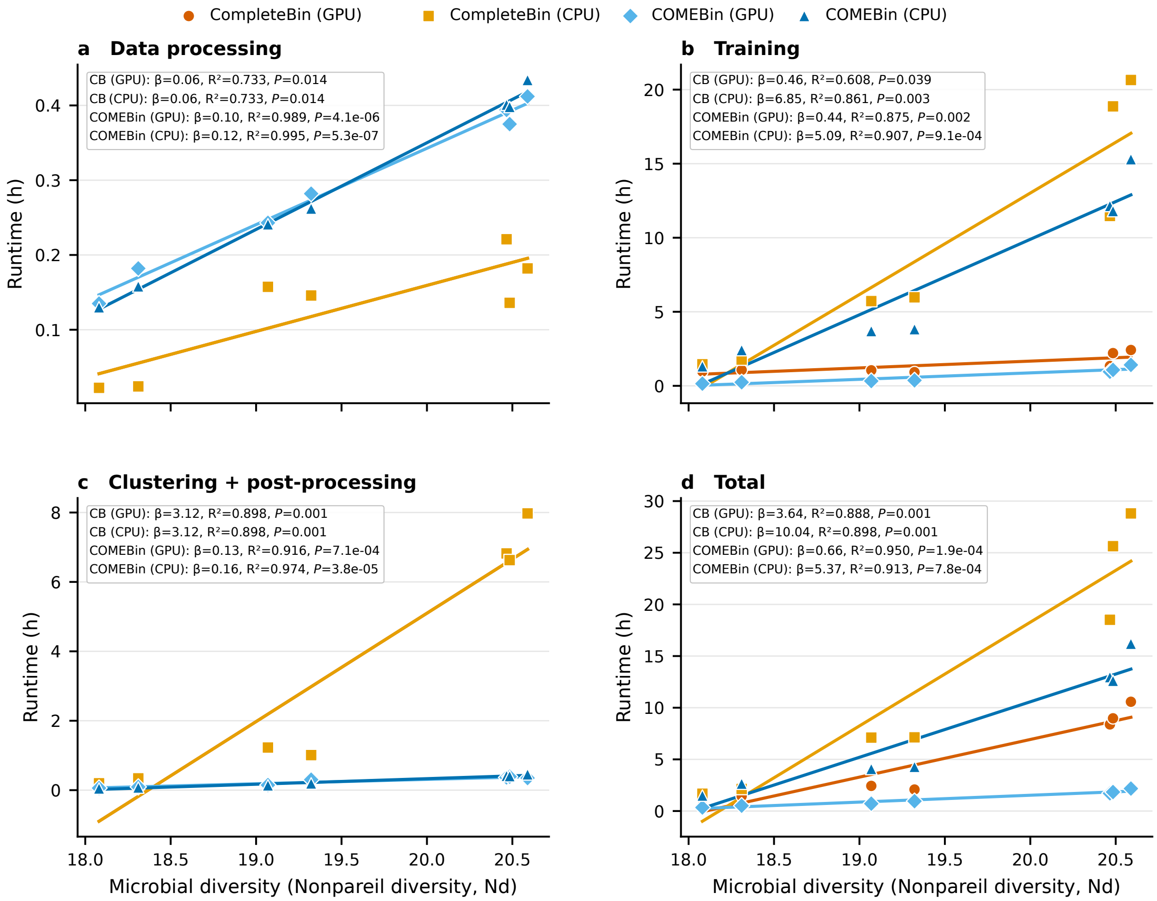


**Supplementary Figure 4: Association between microbial diversity and runtime.** (**a**–**d**) Wall-clock runtime as a function of Nonpareil diversity (Nd) for (**a**) data processing, (**b**) training, (**c**) clustering and post-processing and (**d**) the complete workflow, respectively. Points represent seven CAMI II samples, and lines show linear regressions fitted separately for each execution mode. *P* values are two-sided tests of the null hypothesis that the regression slope equals zero; no adjustment was made for multiple comparisons.

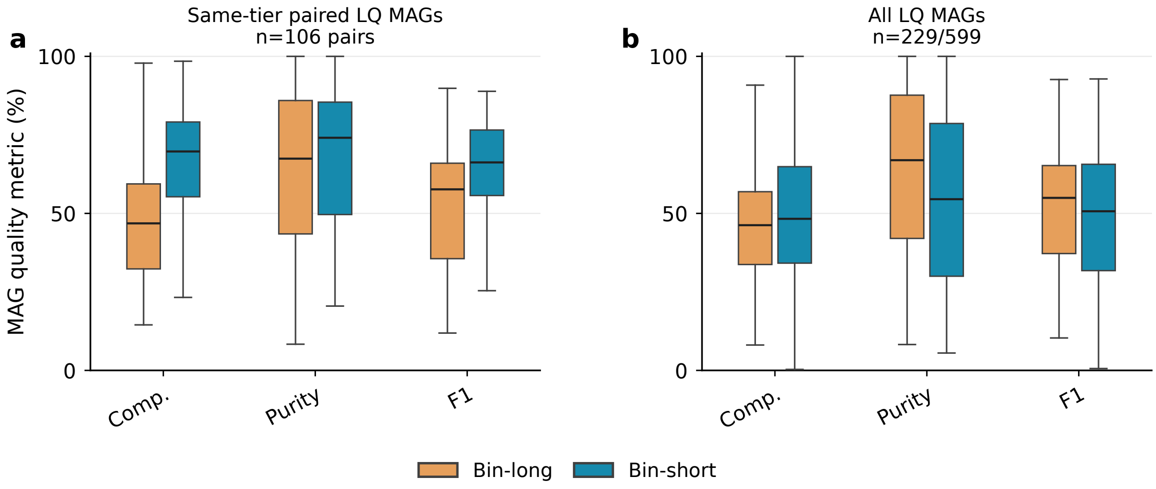


**Supplementary Figure 5: Quality of low-quality MAGs recovered with and without short contigs.** (**a**) Completeness, purity and F1 score for 106 paired low-quality MAGs recovered by bin-long (>2,500 bp) and bin-short (≥850 bp). (**b**) Corresponding distributions for all low-quality MAGs recovered by bin-long ($n = 229$) and bin-short ($n = 599$).

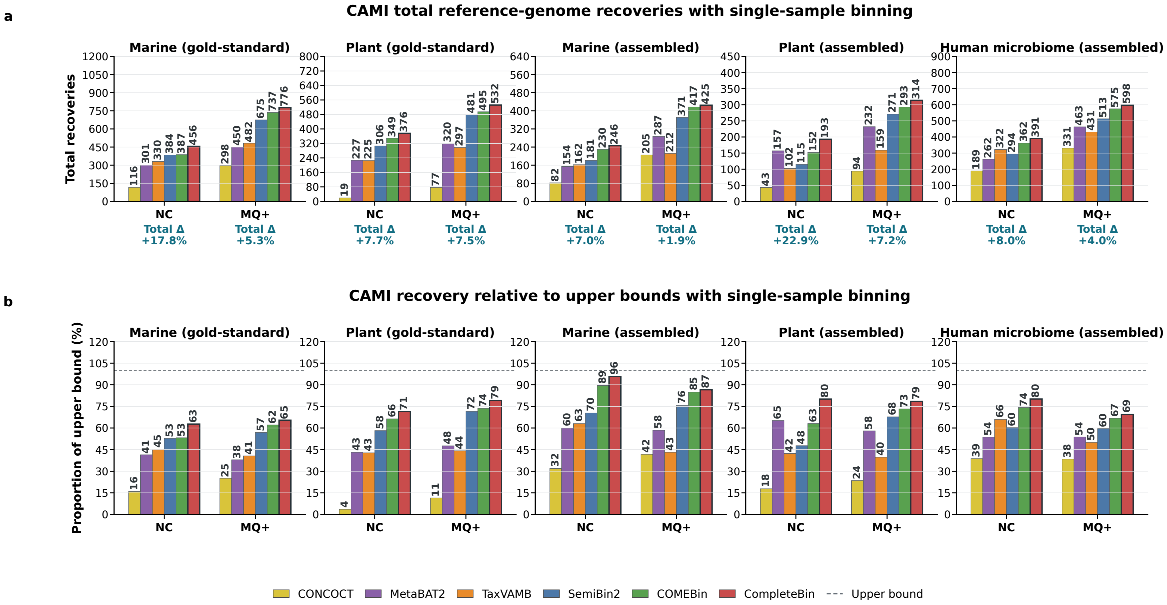


**Supplementary Figure 6: Total reference-genome recovery and recovery relative to assembly-derived upper bounds in CAMI II single-sample binning.** (**a**) Total numbers of reference genomes recovered as near-complete (NC) or medium-quality-or-higher (MQ+) MAGs by six binners across five dataset groups. (**b**) Corresponding recoveries expressed as percentages of the assembly-derived upper bounds; dashed lines indicate the upper bounds. Total Δ values show the relative improvement of CompleteBin over the second-best method.

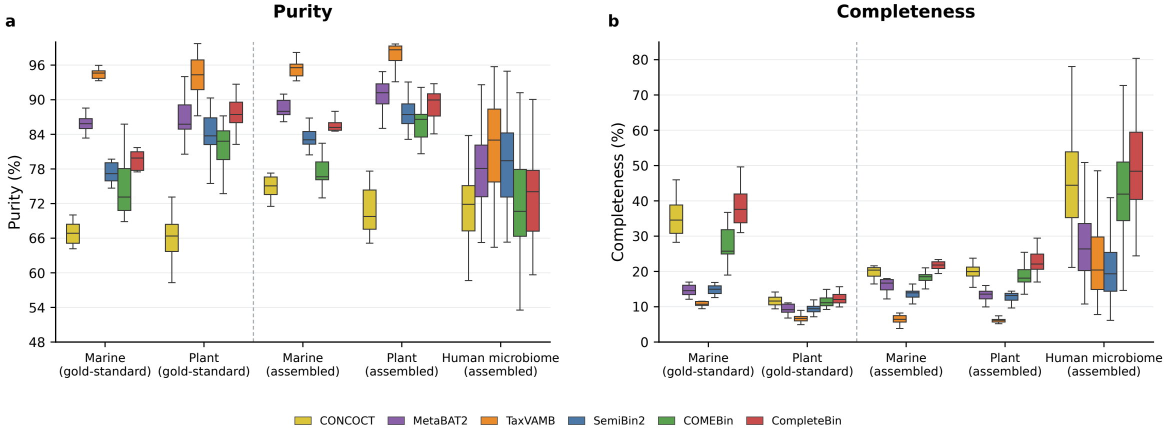


**Supplementary Figure 7: Single-sample binning quality across CAMI II datasets.** Distributions of purity (**a**) and completeness (**b**) for six binning methods on Marine and Plant gold-standard and metagenome assemblies and human-microbiome metagenome assemblies. Colours denote binning methods.

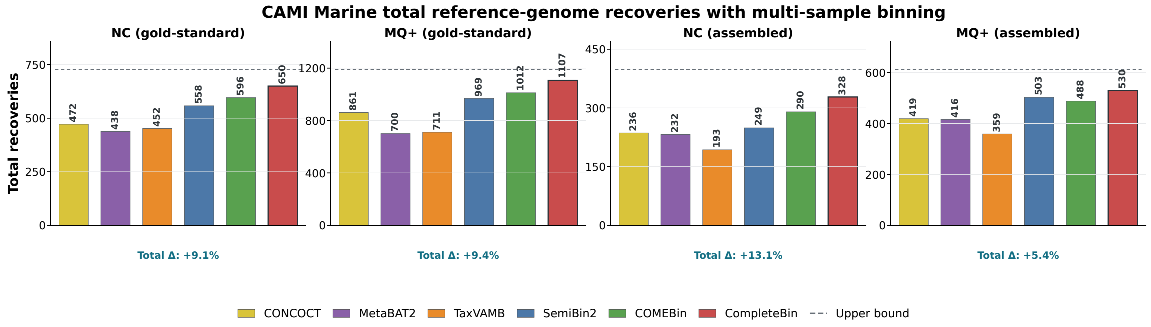


**Supplementary Figure 8: Total reference-genome recovery in CAMI II Marine multi-sample binning.** Total numbers of reference genomes recovered as near-complete (NC) or medium-quality-or-higher (MQ+) MAGs by six binners using gold-standard and metagenome assemblies. Dashed lines indicate the assembly-derived upper bounds, and Total Δ values show the relative improvement of CompleteBin over the second-best method.

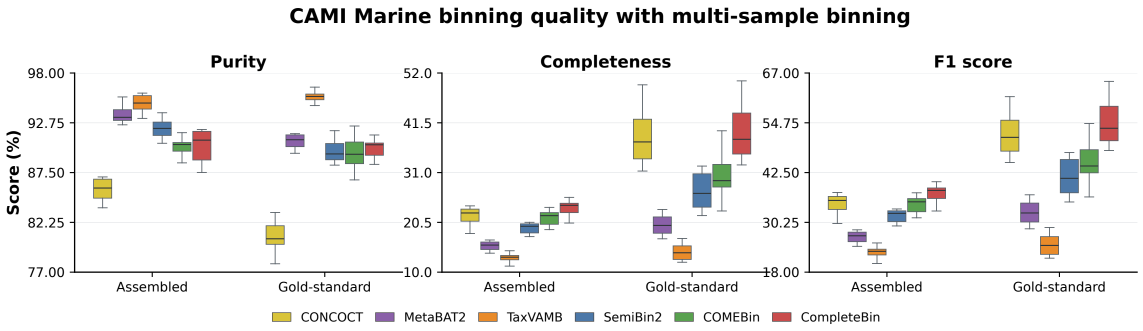


**Supplementary Figure 9: Binning quality for CAMI II Marine multi-sample binning.** Distributions of purity, completeness and F1 score across ten samples for six binning methods using metagenome and gold-standard assemblies. Colours denote binning methods.

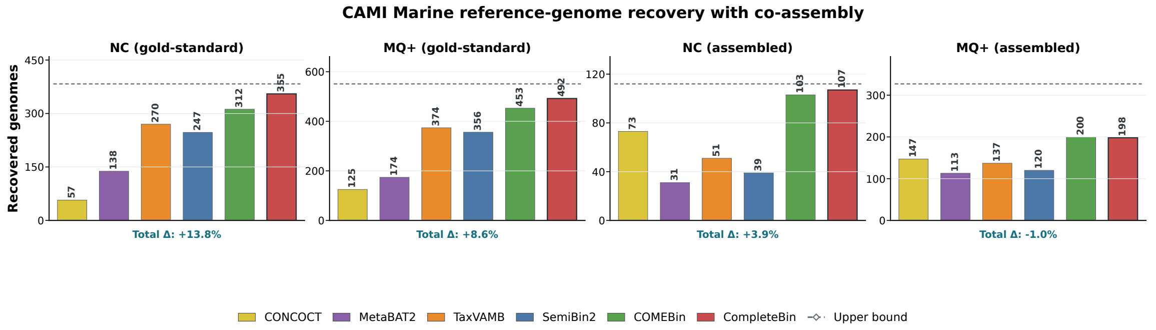


**Supplementary Figure 10: Reference-genome recovery in CAMI II Marine co-assembly binning.** Numbers of reference genomes recovered as near-complete (NC) or medium-quality-or-higher (MQ+) MAGs by six binners using gold-standard and metagenome co-assemblies. Dashed lines indicate the assembly-derived upper bounds, and Total Δ values show the relative difference between CompleteBin and the best-performing alternative.

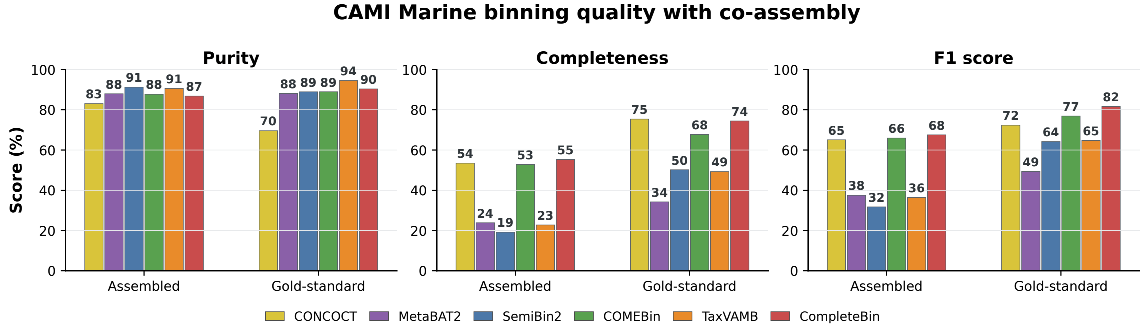


**Supplementary Figure 11: Binning quality on CAMI II Marine co-assemblies.** Purity, completeness and F1 score achieved by six binning methods using the metagenome and gold-standard co-assemblies. Values above the bars indicate scores (%), and colours denote binning methods.

### References


[1] Duncan, A. *et al.* Metagenome-assembled genomes of phytoplankton microbiomes from the arctic and atlantic oceans. *Microbiome* **10**, 67 (2022).

[2] Rodriguez-r, L. M. & Konstantinidis, K. T. Nonpareil: a redundancy-based approach to assess the level of coverage in metagenomic datasets. *Bioinformatics* **30**, 629–635 (2014).

[3] LJPvd, M. & Hinton, G. Visualizing high-dimensional data using t-sne. *J Mach Learn Res* **9**, 9 (2008).
